# Functional genomic analysis of non-canonical DNA regulatory elements of the aryl hydrocarbon receptor

**DOI:** 10.1101/2023.05.01.538985

**Authors:** Shayan Shahriar, Tajhal D Patel, Manjula Nakka, Sandra L Grimm, Cristian Coarfa, Daniel A Gorelick

## Abstract

The aryl hydrocarbon receptor (AHR) is a ligand-dependent transcription factor activated by environmental toxicants like halogenated and polycyclic aromatic hydrocarbons, which then binds to DNA and regulates gene expression. AHR is implicated in numerous physiological processes, including liver and immune function, cell cycle control, oncogenesis, and metabolism. Traditionally, AHR binds a consensus DNA sequence (GCGTG), the xenobiotic response element (XRE), recruits coregulators, and modulates gene expression. Yet, recent evidence suggests AHR can also regulate gene expression via a non-consensus sequence (GGGA), termed the non-consensus XRE (NC-XRE). The prevalence and functional significance of NC-XRE motifs in the genome have remained unclear. While ChIP and reporter studies hinted at AHR-NC-XRE interactions, direct evidence for transcriptional regulation in a native context was lacking. In this study, we analyzed AHR binding to NC-XRE sequences genome-wide in mouse liver, integrating ChIP-seq and RNA-seq data to identify candidate AHR target genes containing NC-XRE motifs in their regulatory regions. We found NC-XRE motifs in 82% of AHR-bound DNA, significantly enriched compared to random regions, and present in promoters and enhancers of AHR targets. Functional genomics on the Serpine1 gene revealed that deleting NC-XRE motifs reduced TCDD-induced Serpine1 upregulation, demonstrating direct regulation. These findings provide the first direct evidence for AHR-mediated regulation via NC-XRE in a natural genomic context, advancing our understanding of AHR-bound DNA and its impact on gene expression and physiological relevance.

## INTRODUCTION

The toxic effects of secondhand cigarette smoke, fossil fuels, and byproducts of industrial combustion are caused by halogenated and polycyclic aromatic hydrocarbons such as TCDD. Understanding the signaling pathways by which such compounds cause toxicity is crucial for predicting tolerable exposure levels and for reversing the effects of adverse exposure. Aromatic hydrocarbons activate aryl hydrocarbon receptors (AHR), a ligand-dependent transcription factor. Upon activation, AHR translocates to the nucleus, where it recruits protein coregulators and directly regulates gene expression (Denison *et al*., 2011). AHR forms a dimer with the aryl hydrocarbon receptor nuclear translocator protein (ARNT, also known as HIF1β) to regulate transcription (Reyes *et al*., 1992; Hankinson, 2005). The AHR-ARNT complex binds to a consensus DNA sequence (GCGTG) termed the xenobiotic response element (XRE) (Denison *et al*., 1988; Yao and Denison, 1992).

Recent evidence complicates these findings. AHR may regulate gene expression by binding to non-consensus XRE sequences. AHR was shown to physically interact with RelB, a member of the NF-κB family, and bind to a distinct RelB/AHR responsive element (RelBAhRE) within the IL-8 promoter. Furthermore, TCDD induced a time-dependent recruitment of AHR to the RelBAhRE site (nucleotides GGGTGCAT) (Vogel *et al*., 2007). A chromatin immunoprecipitation study found that approximately half of TCDD-AHR complexes in mouse liver were bound to DNA lacking a consensus XRE (Dere *et al*., 2011). Exploring the genome more closely, Huang and colleagues identified a non-consensus DNA sequence to which AHR binds (Huang and Elferink, 2012). Chromatin immunoprecipitation (ChIP), gel shift (electrophoretic mobility shift assay) and reporter assays support the hypothesis that AHR can bind to a non-consensus XRE DNA sequence characterized by the nucleotides GGGA (referred to as the NC-XRE motif) (Jackson *et al*., 2014; Joshi *et al*., 2015; Wilson *et al*., 2013; Huang and Elferink, 2012), though this has not been functionally tested in a natural genomic context. Here, we sought to analyze AHR binding to non-canonical DNA elements on a genome-wide scale and, in the case of a single target gene, directly test whether NC-XRE motifs are required for AHR-dependent target gene expression.

## RESULTS

### Frequency and distribution of non-canonical AHR binding motifs

To determine the genome-wide distribution of AHR binding to DNA following TCDD exposure, we analyzed published AHR ChIP-seq data from livers of C57BL/6 male and female mice (28 days old) following 2 hour exposure to 30 μg/kg TCDD (GEO accession numbers GSE97634, GSE97636) (Fader *et al*., 2017, 2019). This data was previously assayed for AHR binding to consensus XRE DNA [GCGTG] but was not interrogated for AHR binding to non-consensus DNA such as NC-XRE [GGGA] or RelBAhRE [GGGTGCAT]. Given that the XRE, NC-XRE, and RelBAhRE motifs differ in length, which influences their frequency of occurrence across the genome (shorter motifs being more common), we assessed the likelihood of finding XRE, NC-XRE or RelBAhRE motifs in AHR ChIP peaks versus in random genomic intervals matched in size and chromosome distribution to observed peaks. Both XRE and NC-XRE motifs were significantly enriched in AHR-bound regions, whereas RelBAhRE was not (Fig. 1A). These random regions were scanned with the same motif definitions as the observed peaks, ensuring that motif length–dependent differences in background occurrence were directly incorporated into the enrichment test.

**Figure 1.**
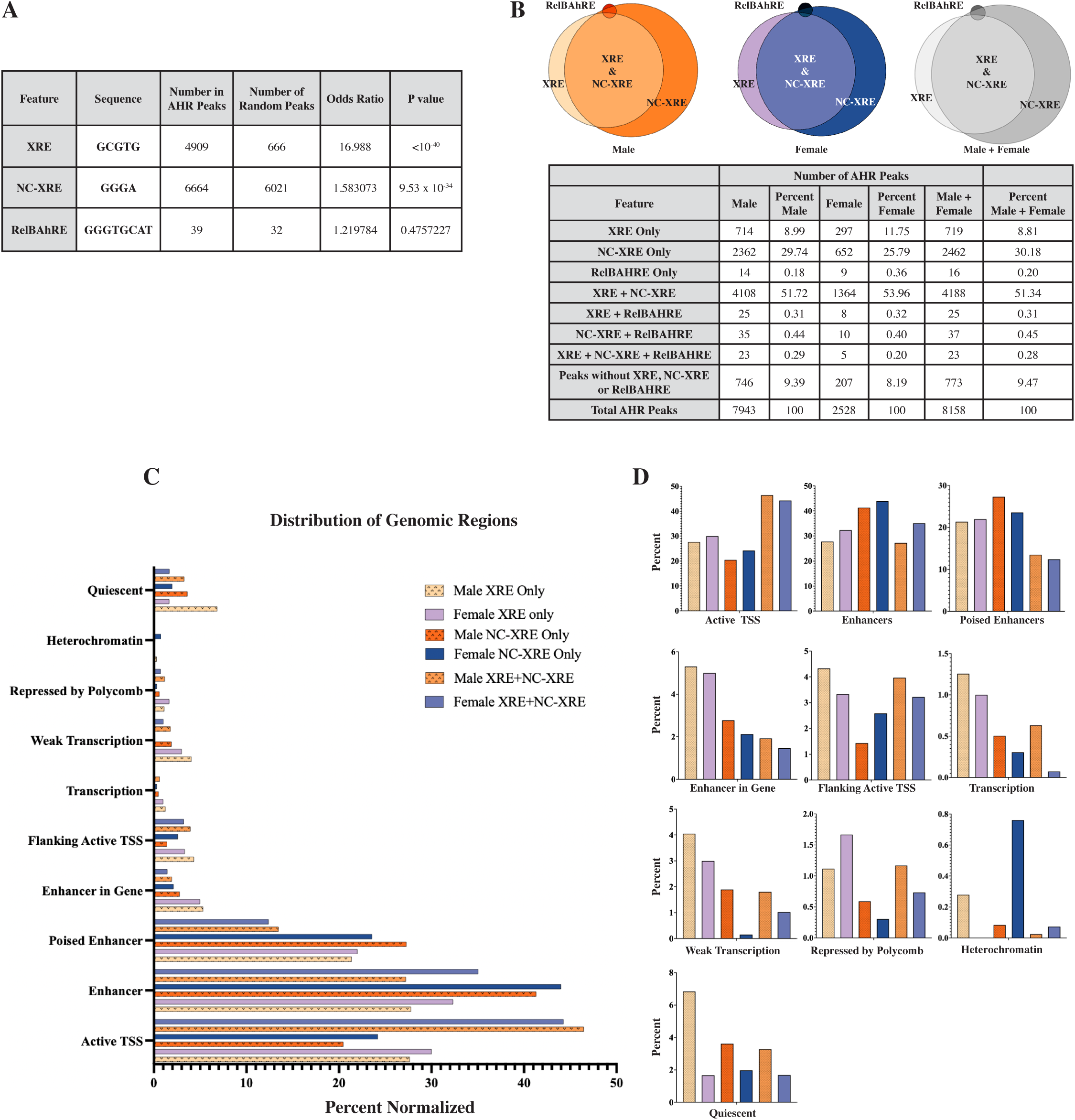
Enrichment and Distribution of NC-XRE, XRE, and RelBAhRE motifs in AHR ChIP-seq Peaks. Following TCDD exposure, we analyzed mouse liver AHR ChIP-seq peaks from 5 male and 5 female mice for the presence of known AHR motifs: canonical xenobiotic response element (XRE), non-canonical xenobiotic response element (NC-XRE), and RelB AHR response element (RelBAHRE). (A) Likelihood of finding XRE, NC-XRE, or RelBAHRE motifs in AHR ChIP peaks versus in random genomic regions. The table shows the number of AHR ChIP peaks containing at least one XRE, NC-XRE, or RelBAHRE motif. Odds ratios and p-values derived from Fisher’s exact test indicate significant enrichment of XRE and NC-XRE motifs in AHR peaks. (B) Proportion of AHR ChIP peaks containing XRE DNA sequence, NC-XRE, XRE + NC-XRE, or none in male and female mice. NC-XRE motifs are more prevalent than XRE motifs in both sexes. (C) Genomic distribution of AHR ChIP-seq peaks mapped to 10 chromatin states defined by ChromHMM using 8 histone modification profiles. The highest percentages of AHR peaks containing XRE and NC-XRE motifs are found in promoter and enhancer regions. (D) Enlarged view of panel C for detailed comparison of chromatin state distributions. This panel highlights the differences in the distribution of AHR peaks in various chromatin states. Overlapping XRE and NC-XRE motifs are slightly more prominent in promoters; NC-XREs without XREs are slightly more enriched in enhancers and poised enhancers; XREs without NC-XREs are more enriched at enhancers in genes and at gene bodies with active or weak transcription.

We also found consistent patterns of AHR-bound motif enrichment in the livers of male and female mice following TCDD exposure, based on AHR ChIP-seq data. In males, 29.74% of AHR peaks contained only NC-XRE motifs, compared to 25.79% in females. Peaks containing only XRE motifs were slightly more frequent in females (11.75%) than in males (8.99%). The majority of peaks—51.72% in males and 53.96% in females— contained both XRE and NC-XRE motifs, indicating frequent co-occurrence. Despite these minor differences, the overall distribution of motif combinations was remarkably similar between sexes. RelBAhRE motifs were rare in both groups, appearing in fewer than 1% of peaks (Fig. 1B and Supplementary Fig. 1A). These findings indicate that AHR binding to canonical and non-canonical DNA motifs occurs at comparable frequencies in male and female mouse livers, supporting a robust and sex-independent pattern of AHR-DNA interactions in response to TCDD. They also highlight the substantial role of NC-XRE motifs in mediating AHR binding. Because of the low number of RelBAhRE in AHR peaks, we only focused on the XRE and NC-XRE motifs in the subsequent studies.

We next examined how AHR binding sites containing XRE and NC-XRE motifs are distributed across the genome. To do this, we analyzed AHR ChIP-seq peaks in adult mouse liver (GEO accession numbers GSE97634, GSE97636) and overlaid them with genomic regions annotated using a 10-state ChromHMM model (van der Velde *et al*., 2021). We generated this model using eight histone modifications profiled via publicly available ChIP-seq datasets (GEO accession numbers: GSM1000140, GSM918718, GSM1000150, GSM1000151, GSM1000152, GSM1000153, GSM769014, GSM769015, GSM751035) (Mouse ENCODE Consortium *et al*., 2012; Tennant *et al*., 2013). These included histone marks associated with active promoters (H3K4me3, H3K9ac), enhancers (H3K4me1, H3K27ac), gene bodies and active transcription (H3K36me3), polycomb-mediated repression (H3K27me3), and heterochromatin (H3K9me3) (Supplementary Figure 2E).

We found that AHR binding was enriched in active promoters (e.g. active TSS) and enhancers (Figure 1C, 1D). Specifically, AHR peaks containing both XRE and NC-XRE motifs were predominantly located in active transcription start sites (TSS) and active enhancers. On the other hand, most NC-XRE-only peaks were found in the enhancer regions, including both active and poised enhancers. Active enhancers have high chromatin accessibility and confer strong enhancer activity whereas poised enhancers are marked by intermediate levels of enhancer-associated histone modifications and are thought to be primed for activation. We also noticed that AHR binding was rare in repressive chromatin states, such as those enriched for H3K27me3 (polycomb repression) or H3K9me3 (heterochromatin), as well as in the quiescent regions lacking significant histone modification signals. These patterns were consistent across male and female samples, supporting a sex-independent mechanism of AHR-DNA interaction (Figure 1C, 1D). Notably, promoter- and enhancer-associated regions comprise only ∼5.3% of the mouse genome, whereas repressive and quiescent regions account for approximately 84% (van der Velde *et al*., 2021). This highlights a strong preference of AHR for binding within the active regulatory elements, consistent with its function as a transcription factor. Specifically, NC-XRE motifs may play a particularly important role in enhancer-associated AHR binding.

### Repeated NC-XRE motifs associated with AHR ChIP peaks

We searched for NC-XRE motifs present near AHR binding sites (ChIP peak) within 10 kb of a gene body. The 10 kb parameter allows us to include promoter and gene proximal regulatory DNA peaks, important because our analysis of ChIP-seq data suggested that AHR binds NC-XRE at enhancers (Figure 1C, D). Among the differentially expressed genes in TCDD vs vehicle, 520 had AHR binding peaks at XRE, NC-XRE or XRE+NC-XRE motifs within 10 kb of a gene body. 28 of these genes had AHR binding peaks at NC-XRE but not at XRE or XRE+NC-XRE motifs (Supplementary Table 1).

The preceding analysis is for any single instance of XRE or NC-XRE DNA sequence. However, for some transcription factors, repeated DNA binding sites produce a more robust response (Klein-Hitpass *et al*., 1988). To explore the functional importance of repeated runs of XRE or NC-XRE motifs, we sought to examine the association between repeated runs of XRE or NC-XRE motifs and AHR target gene expression.

We defined tandem XRE and NC-XRE motifs based on two parameters: the number of repeats and the maximum allowed spacer length (in base pairs) between adjacent motifs, as illustrated in Figure 2A. The number of repeats refers to the minimum number of XRE or NC-XRE motifs within a given region, while the spacer defines the maximum distance permitted between any two motifs in a repeat run. Using this framework, we analyzed AHR target genes for varying numbers of XRE or NC-XRE repeats and spacer lengths. Notably, the previously characterized AHR target gene *Serpine1* (encodes the protein PAI1) contains five NC-XRE motifs within its putative promoter region. This observation aligns with previous findings that AHR upregulates *Serpine1* via NC-XRE-mediated binding (Huang and Elferink, 2012), prompting us to further investigate the regulatory significance of five-repeat NC-XRE clusters.

**Figure 2.**
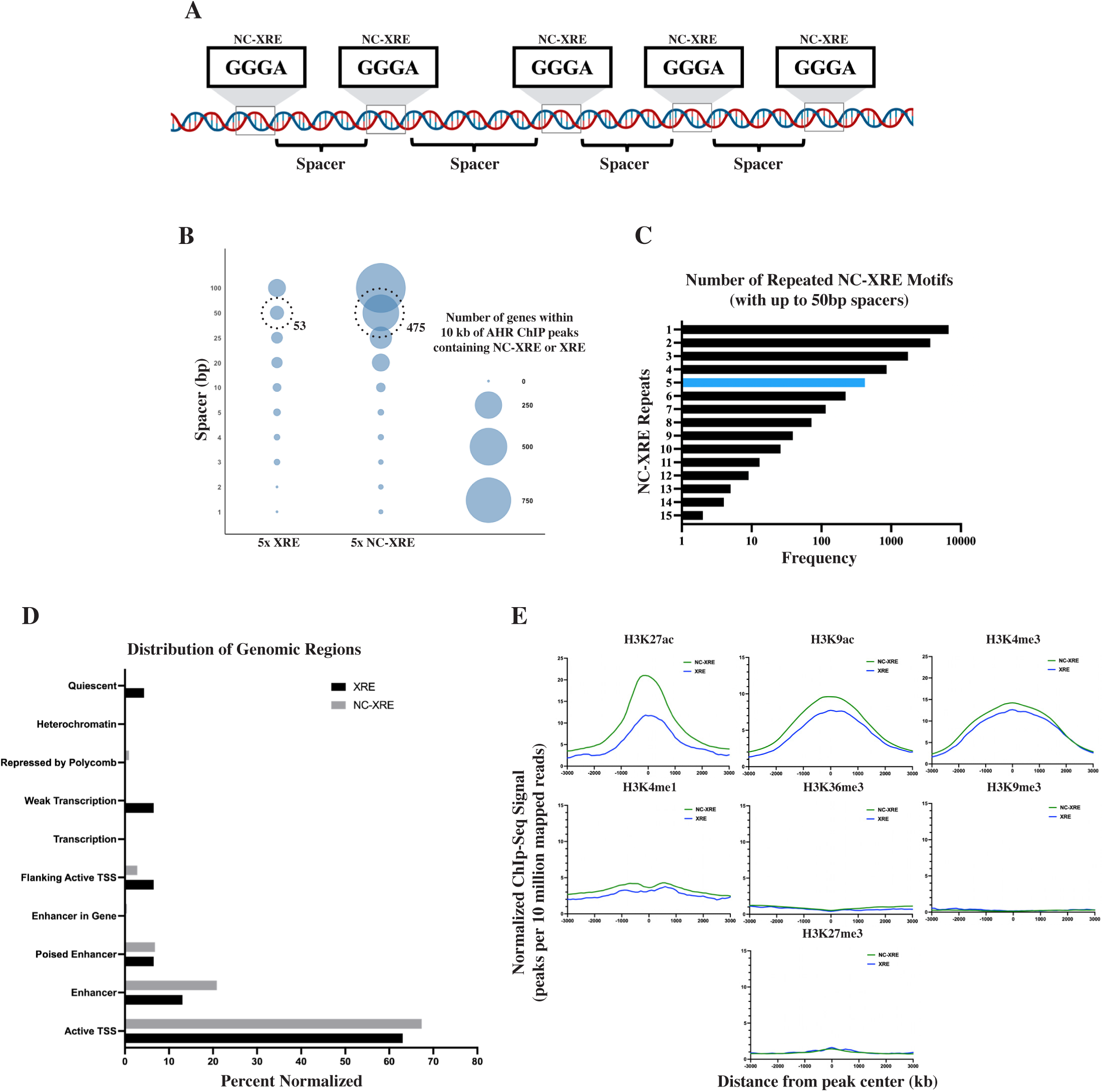
Analysis of NC-XRE Repeats and Their Genomic Context. (A) We categorized NC-XRE DNA motifs based on the number of repeated NC-XRE motifs and the distance separating those motifs (spacers) from each other. For example, 5 motifs separated by 50 basepairs (bp) would indicate 5 NC-XRE motifs where each spacer is up to 50 bp long. (B) Number of genes within 10 kb of AHR ChIP-seq peaks containing either 5 repeats of NC-XRE or XRE motifs separated by spacers ranging from 1 to 100 basepairs (bp). The size of the circles represents the number of AHR target genes. Highlighted circles denote that there are 9-fold more AHR target genes associated with AHR ChIP peaks containing 5 NC-XRE motifs versus 5 XRE motifs separated by 50 bp or less. (C) Frequency of repeated runs of NC-XRE motifs in AHR ChIP peaks. It shows the total number of AHR ChIP peaks containing different numbers of NC-XRE repeats (1-15) with a maximum of 50 bp spacer between any two motifs. (D) Genomic distribution of AHR ChIP-seq peaks containing 5 XRE or NC-XRE motifs separated by 50 bp or less. These are mapped to 10 chromatin states defined by ChromHMM using 8 histone modification states. The highest percentages of AHR peaks containing XRE and NC-XRE motifs are found in promoter and enhancer regions. NC-XRE motifs are prevalent in enhancers and promoters compared to XRE motifs but are absent in gene bodies (transcription, weak transcription, enhancer in gene). (E) Binding strength of indicated histone modifiers at AHR ChIP peaks containing 5 or more NC-XRE or XRE motifs separated by 50 bp or less. AHR-NC-XRE peaks are more associated with activating histone marks (H3K27ac, H3K9ac, H3K4me3, H3K4me1).

Fig 2B shows the number of AHR target genes with AHR peaks containing at least 5 NC-XRE motifs separated by up to 100 basepairs (bp), compared to target genes found near AHR peaks with at least 5 XRE motifs. Notably, we observed a ninefold increase in the number of AHR target genes linked to peaks with five NC-XRE motifs separated by 50 bp or less, compared to those with five XRE motifs under the same spacing constraint. This subset includes *Serpine1*, a well-established AHR target regulated via NC-XRE motifs, as previously discussed (Fig 2B).

Further analysis revealed that while 82% of all AHR peaks contain at least one NC-XRE motif, only 5.2% harbor a cluster of at least five NC-XRE motifs spaced by 50 bp or less (Figure 2C, Supplementary Figure 2A). We next characterized the genomic distribution of these 5x NC-XRE clusters. Of the 426 identified clusters, over 71% were located at or near transcription start sites (TSS), indicating a strong promoter association. An additional 27.8% were found in enhancer regions, with 21% in active enhancers and 6.8% in poised enhancers. These clusters were rarely found in repressed chromatin states (e.g., heterochromatin, polycomb-repressed, or quiescent regions) or within gene bodies associated with transcriptional activity (transcription or weak transcription) (Figure 2D, Supplementary Figures 2C and 2D). Similarly, clusters of five XRE motifs were predominantly located in promoters and enhancers, with a small fraction (6.5%) found within gene bodies. We compared the binding strengths of different histone marks at AHR peaks containing 5x XRE or NC-XRE (50 bp spacer). NC-XREs are enriched at AHR binding sites proximal to transcription start sites (H3K4me3, H3K27ac) and enhancers (H3K27ac) (Fig 2E, Supplementary Fig 2E). Consistent with the broader genomic distribution of NC-XREs, 5X NC-XRE clusters were enriched in enhancer regions compared to 5x XRE clusters. This also suggests that NC-XREs may be more potent when they occur in tandem repeats.

### Putative AHR-NC-XRE target genes and pathways

The preceding analysis examined the presence of NC-XRE motifs in all genomic regions where AHR was bound to DNA. However, transcription factors like AHR may bind DNA without necessarily affecting gene expression. Therefore, to characterize functional AHR binding events, we sought to identify AHR ChIP-seq peaks associated with AHR target genes. For this, we integrated the AHR ChIP-seq data with published RNA-seq data from mouse liver (Nault *et al*., 2018, 2015; Fader *et al*., 2017). To maximize identification of AHR target genes, we included RNA-seq data from mouse livers exposed to 30µg/kg of TCDD, a potent AHR ligand, across multiple durations (2hr, 4hr, 8hr, 12hr, 24hr) of exposure (Tables 1, 2). We defined AHR target genes by differential expression in TCDD-treated mouse liver at 2 hours vs. DMSO, fold change ≥1.5, FDR < 5%. We focused our analysis on the 2-hour timepoint because this is when direct AHR target gene regulation is most prominent, before the onset of the secondary effects. As illustrated in Supplementary Figure 5C, gene set enrichment analysis (discussed in detail later) of AHR-NC-XRE-regulated pathways shows a clear enrichment at 2 hours of TCDD exposure, followed by a decline at later time points. This pattern is consistent with the biology of AHR signaling: TCDD rapidly induces AHR activity and upregulates its direct targets within 2 hours, but by 4 hours and beyond, negative feedback suppresses direct AHR activity. This leads to reduced enrichment of direct AHR target pathways and an increasing contribution of secondary or indirect effects, including pathways regulated by downstream AHR targets.

**Table 1:** ChIP-Seq Datasets Analyzed in This Study.

| GEO Series | PMID | GEO Samples | Target | Sex | Strain | Tissue | Age at Assay | Conditions |
| --- | --- | --- | --- | --- | --- | --- | --- | --- |
| GSE97634 | 28213091<br>31015483 | GSM2573805<br>GSM2573806<br>GSM2573807<br>GSM2573808<br>GSM2573809 | AHR | Male | C57BL/6 | Liver | Adult<br>28 Days | Exposed to 30 ug/kg TCDD via oral gavage for 2 hours, n = 5 mice |
| GSE97636 | 26582802 | GSM2574037<br>GSM2574038<br>GSM2574039<br>GSM2574040<br>GSM2574041 | AHR | Female | C57BL/6 | Liver | Adult<br>28 Days | Exposed to 30 ug/kg TCDD via oral gavage for 2 hours, n = 5 mice |
| GSE31039 | 22889292 | GSM1000140 | H3K27ac | Male | C57BL/6 | Liver | Adult 8 Weeks | No treatment |
|  |  | GSM1000150 | H3K27me3 | Male | C57BL/6 | Liver | Adult 8 Weeks |  |
|  |  | GSM1000151 | H3K36me3 | Male | C57BL/6 | Liver | Adult 8 Weeks |  |
|  |  | GSM1000152 | H3K79me2 | Male | C57BL/6 | Liver | Adult 8 Weeks |  |
|  |  | GSM1000153 | H3K9ac | Male | C57BL/6 | Liver | Adult 8 Weeks |  |
|  |  | GSM769014 | H3K4me3 | Male | C57BL/6 | Liver | Adult 8 Weeks |  |
|  |  | GSM769015 | H3K4me1 | Male | C57BL/6 | Liver | Adult 8 Weeks |  |
| GSE36027 | 22889292 | GSM918718 | Control | Male | C57BL/6 | Liver | Adult 8 Weeks | No treatment |
| GSE30298 | 23238790 | GSM751035 | H3K9me3 | N/A | C57BL/6 | Liver | Adult<br>8-10 Weeks | No treatment |

**Table 2:**
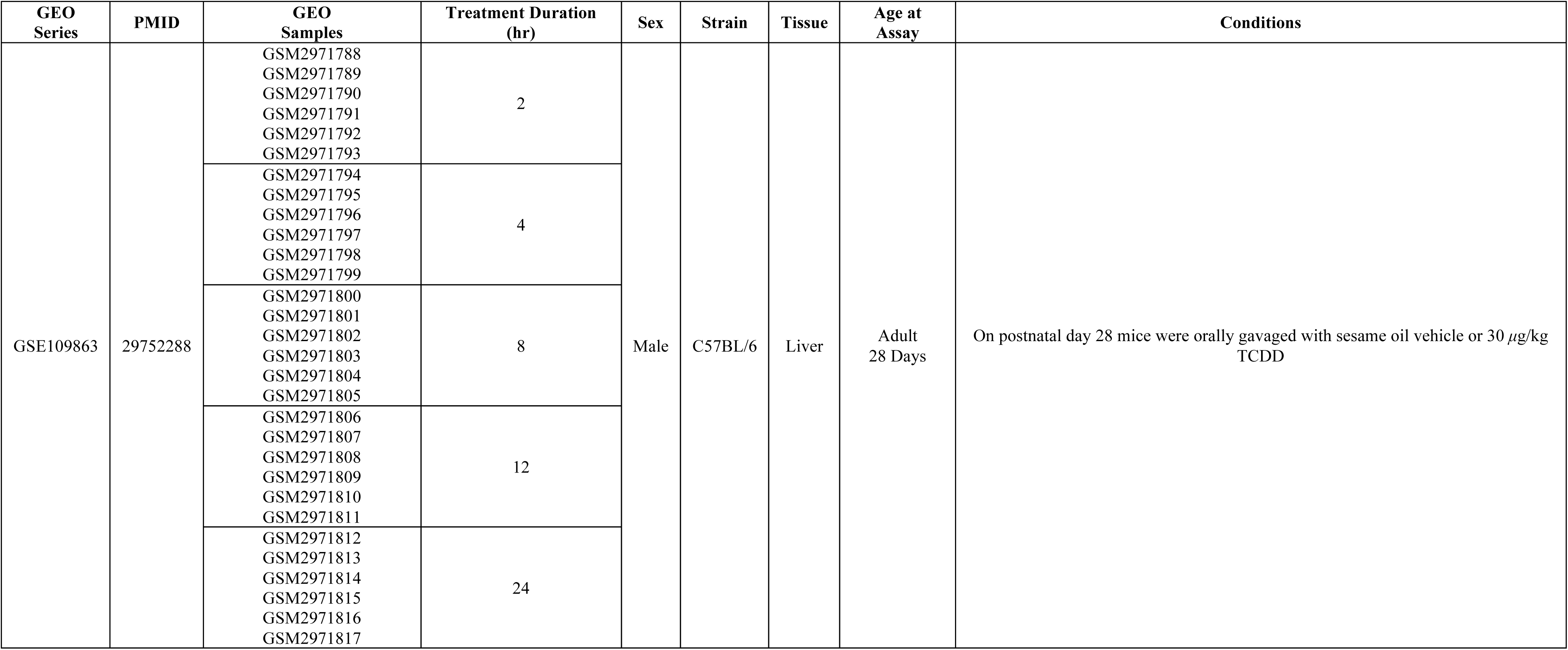
RNA-Seq Datasets Analyzed in This Study.

We examined whether AHR-dependent differential gene expression correlated with the number of NC-XRE motifs by plotting the fold changes of all upregulated AHR target genes after 2 and 4 hours of TCDD exposure against the number of NC-XREs within 10 kb of each gene body, followed by linear regression analysis (Supplementary Figure 5A, 5B). No significant correlation was observed. Given that the known AHR-NC-XRE target *Serpine1* harbors five repeated NC-XRE motifs in its putative promoter, we next focused on AHR ChIP-seq peaks containing at least five NC-XRE motifs spaced ≤50 base pairs apart to identify additional putative AHR targets. Using this approach, we identified such NC-XRE clusters near 28 AHR target genes, among which, 22 genes were upregulated in response to TCDD exposure. These differentially upregulated genes with 5 or more NC-XRE motifs are shown in Fig 3A. As validation, *Serpine1* was among these genes, consistent with prior findings that AHR upregulates *Serpine1* via NC-XRE DNA (Huang and Elferink, 2012). AHR binding was also detected at NC-XRE motifs near genes like *Tiparp*, another known AHR target gene induced by TCDD (Ma *et al*., 2001) and reported to suppress hepatic gluconeogenesis (Diani-Moore *et al*., 2010). These findings suggest that NC-XRE motifs may contribute to the regulation of *Tiparp* expression, pointing to a potential novel mechanism underlying TCDD-induced hepatotoxicity.

**Figure 3.**
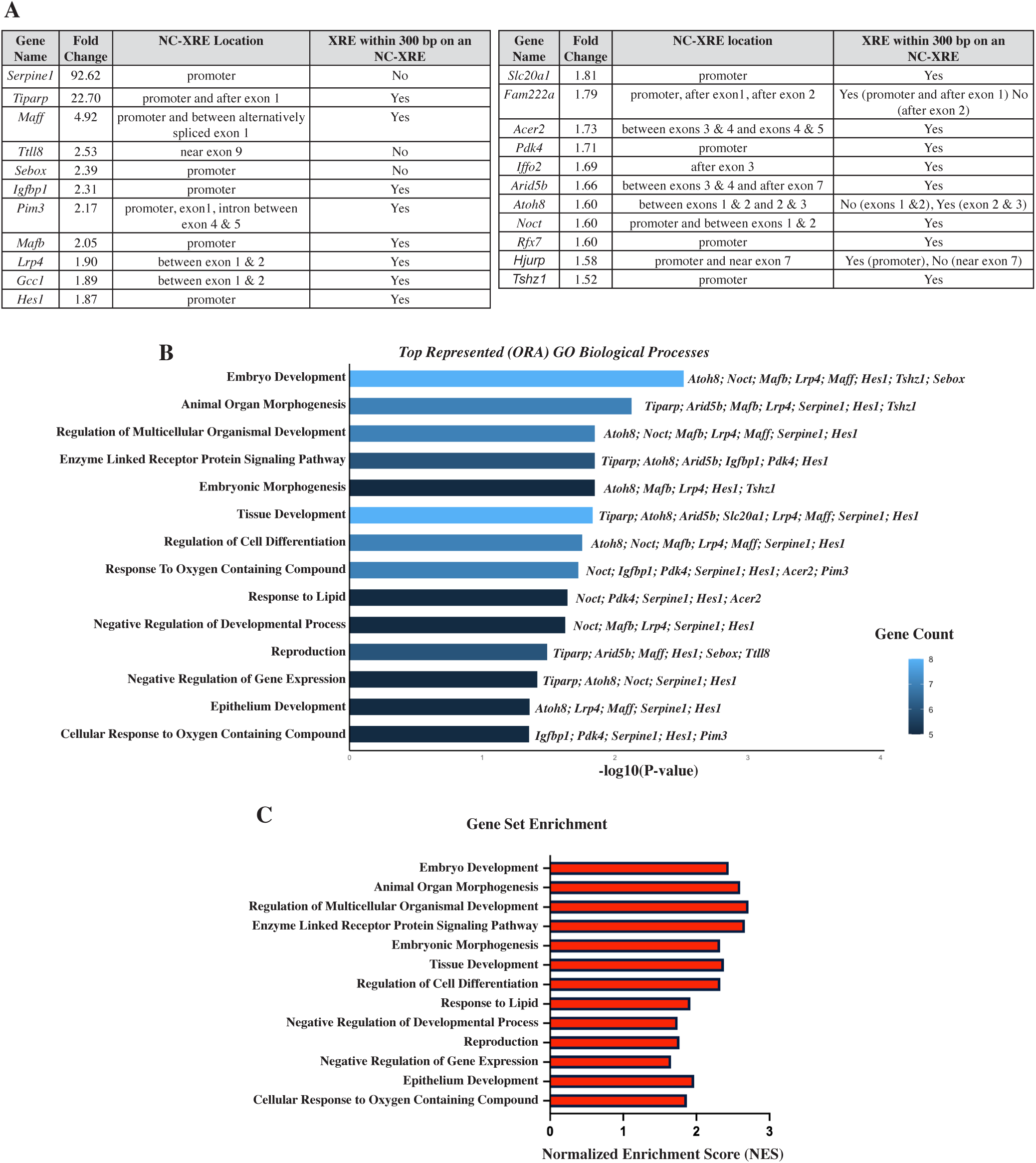
Analysis of AHR target genes. (A) Upregulated AHR target genes (fold change TCDD vs vehicle, 2-hour exposure, mouse liver) containing AHR ChIP peaks with 5 or more NC-XRE motifs separated by 50 base pairs or less. AHR target genes are defined as differentially expressed genes (RNA-seq, fold difference in TCDD vs vehicle > 1.5x, FDR < 5%) containing AHR ChIP peaks within 10 kb of a gene body. (B) Over-Representation Analysis (ORA) of Gene Ontology Biological Processes (GOBP) using the AHR target genes from panel A. Enriched pathways were ranked by their statistical significance (P-value) along with the number of hit genes and their names. For example, the embryo development GOBP pathway contains 8 upregulated genes (Atoh8, Notch1, Mafb, Lrp4, Maff, Hes1, Tsgz1, Sebox) that have 5 repeats of NC-XRE motifs with a maximum of 50 bp spacers between them. (C) Gene Set Enrichment Analysis (GSEA) displaying normalized enrichment scores (NES) for mouse liver global RNA-seq (TCDD vs vehicle) ranked genes. Only the pathways identified in the ORA (panel B) are shown here. All except the ‘response to oxygen-containing compound’ pathway are enriched in the TCDD-exposed mouse liver.

To further explore the biological relevance of the 22 upregulated AHR target genes containing 5X NC-XRE motifs (50 bp spacers), we performed an over-representation analysis (ORA) to identify significantly related Gene Ontology (GO) biological processes (FDR < 5%). This analysis revealed 14 pathways (Figure 3B). However, many of these pathways are broad and involve large gene sets—for example, the GO term “embryo development” includes over 1,000 genes. To contextualize these findings, we conducted a gene set enrichment analysis (GSEA) using a global ranked list of genes based on differential expression in mouse liver following 2-hour TCDD treatment compared to DMSO. This approach allowed us to assess whether the identified pathways were enriched across the entire transcriptome. 13 out of the 14 pathways identified in the ORA were also significantly enriched in the GSEA (FDR < 25%) (Figure 3C).

The enriched pathways span a range of biological functions, including development, morphogenesis, oxygen response, and enzymatic activity—processes that align with known physiological roles of AHR. These findings reveal a compelling link between AHR-NC-XRE target gene expression and key biological pathways, suggesting that AHR may regulate these biological processes, at least in part, through NC-XRE-mediated transcriptional control.

### Putative transcription factors regulated via NC-XRE motifs

Transcription factors bind regulatory DNA in a complex composed of multiple different transcription factors. For example, AHR is associated with the transcription factor estrogen receptor alpha at regulatory DNA (Beischlag and Perdew, 2005; Klinge *et al*., 2000; Madak-Erdogan and Katzenellenbogen, 2012). Another transcription factor, ARNT, is a co-regulator of AHR (Reyes *et al*., 1992; Lo and Matthews, 2012).

Previous studies have focused on transcription factor binding at XRE motifs. To identify transcription factors that might form a complex with AHR at NC-XRE motifs, we identified known transcription factor DNA binding sites flanking NC-XRE motifs. We focused on AHR ChIP peaks containing at least 2 NC-XRE motifs within 25 basepairs of each other. This threshold was chosen to maximize the detection of potential transcription factors adjacent to NC-XRE bound AHR peaks. It includes all high-density 5x NC-XRE (50 bp spacer) clusters and captures additional AHR-NC-XRE-adjacent regions, making it more permissive for potential transcription factor screening. The top 12 most frequently occurring known transcription factor DNA binding sites are shown in Figure 4A.

**Figure 4.**
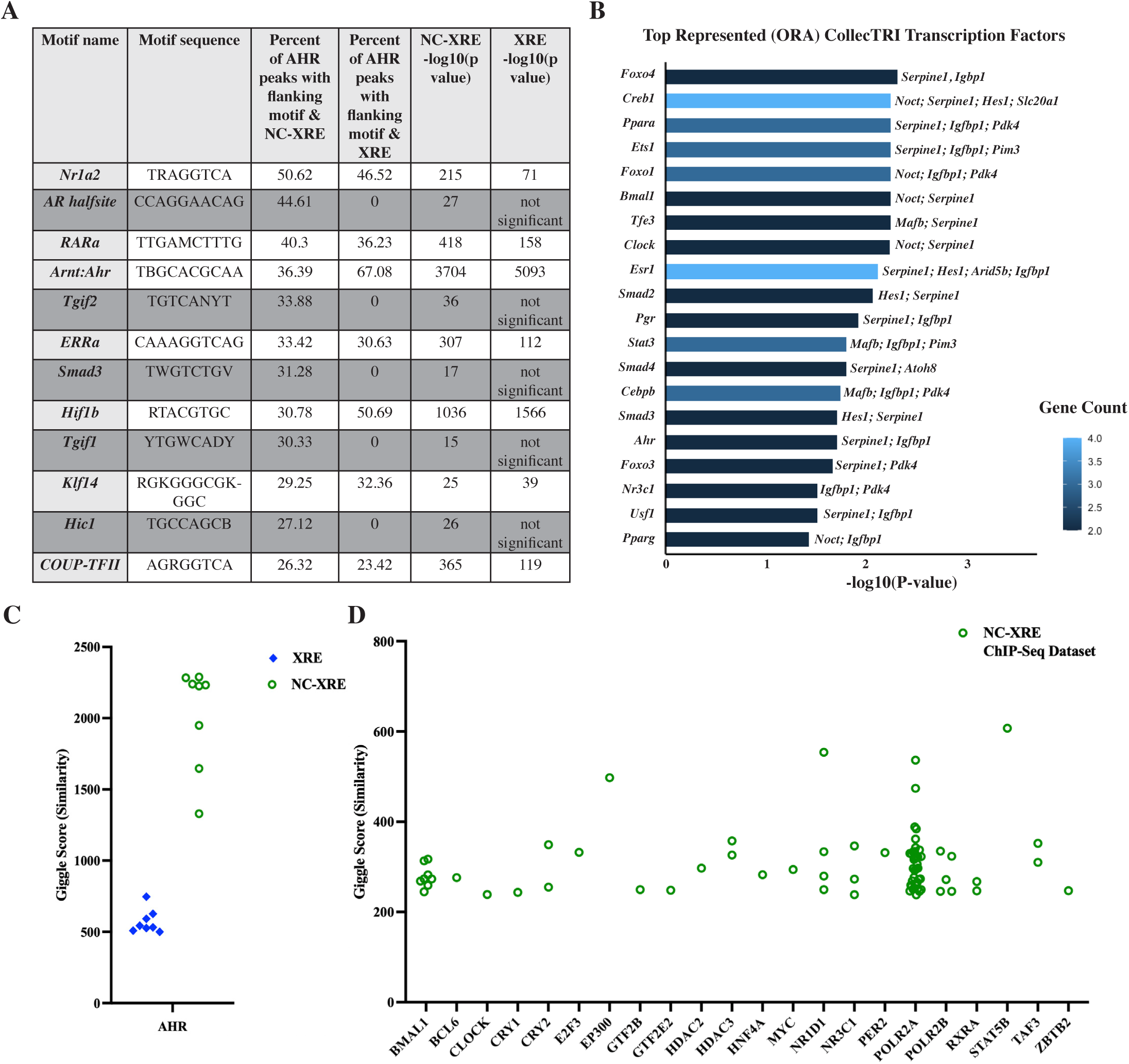
Putative transcription factors acting with AHR at NC-XRE DNA. (A) Top 12 most frequently occurring known transcription factor motifs in the 50 basepairs flanking repeated NC-XRE and XRE motifs (2 or more motifs within 25 basepairs of each other) in genomic regions where AHR binds. Highlighted motifs are enriched flanking NC-XRE but not XRE. Significance defined as p<10^-11^. (B) Over-Representation Analysis (ORA) using the CollecTRI comprehensive gene regulatory network to identify transcription factors known to upregulate AHR targets (described in Figure 3A), where the AHR target genes are associated with 5 NC-XRE repeats separated by 50 bp or less. Top represented transcription factors ranked by their statistical significance (-log10(P-value)). For example, FOXO4 has two hits (*Serpine1, Igfbp1*), indicating that the *Serpine1* and *Igfbp1* genes are (1) targets of the FOXO4 transcription factor protein, (2) are targets of AHR (RNAseq from TCDD in mouse liver), and (3) contain 5 NC-XRE repeats with 50 bp spacers in AHR ChIP peaks within 10 kb of the gene body (ChIP-seq from TCDD in mouse liver). (C, D) Giggle scores of AHR peaks with 5 or more XRE or NC-XRE repeats with up to 50 bp spacers compared to experimentally determined transcription factor peaks from mouse liver or liver cell ChIP-seq. Each point represents how strongly XRE or NC-XRE containing AHR binding sites overlap with the known transcription factor binding sites. A higher giggle score indicates a higher likelihood of these transcription factors co-occupying the AHR peaks. For example, DNA sequence to which STAT5B binds is enriched in AHR ChIP peaks containing NC-XRE (5 or more repeats with 50 bp spacers). Positive control (C) shows high enrichment of AHR DNA binding motifs within AHR ChIP peaks.

As a complementary approach to identify transcription factors that might form a complex with AHR at NC-XRE motifs, we took the 22 upregulated genes mentioned in Figure 3A and asked whether any combination of these genes is known to be targets of additional transcription factors, besides AHR. We queried the CollecTRI database—a curated resource of transcriptional regulators and their experimentally validated targets, commonly used to infer regulon activity (Müller-Dott *et al*., 2023). A regulon is defined as a group of genes regulated by a transcription factor through direct binding to specific regulatory DNA motifs. This analysis revealed 20 significantly overrepresented regulons (FDR < 5%), indicating that multiple transcription factors may form a transcription complex with AHR at NC-XRE DNA (Fig 4B).

To further investigate potential transcription factors that might form a complex with AHR at NC-XRE motifs, we searched for transcription factors that physically bind to the same genomic regions as AHR. We focused on AHR ChIP-seq peaks containing five NC-XRE motifs spaced by 50 base pairs or less. Using the CistromeDB Toolkit, we calculated GIGGLE scores to identify ChIP-seq datasets from the Cistrome Data Browser that significantly overlap with AHR-NC-XRE peak regions (Fig 4C, 4D, Supplementary Fig 4C) (Zheng *et al*., 2019; Mei *et al*., 2017). The GIGGLE score integrates statistical significance (via Fisher’s exact test) and enrichment (odds ratio) to determine how likely the observed overlap is beyond chance (Layer *et al*., 2018). A higher GIGGLE score between two different ChIP-seq datasets (targeting different proteins for the pull down) indicates a strong and statistically meaningful overlap in their peak regions, suggesting that the two different proteins of interest tend to bind in similar genomic locations.

As expected, GIGGLE scores confirmed strong overlap between our AHR-NC-XRE ChIP peak set and existing AHR ChIP-seq datasets, including those originally used to define the 5X NC-XRE peaks. This served as an internal control, validating the sensitivity of our overlap analysis (Figure 4C; Supplementary Figure 4B).

Beyond AHR itself, we identified 23 transcription factors with ChIP-seq peaks from mouse liver or from cultured mouse hepatocytes that significantly overlapped with the 5X NC-XRE (50 bp spacer) AHR peaks (Figure 4D). This suggests that these transcription factors may co-occupy NC-XRE-rich DNA regions alongside AHR. These included factors such as BMAL1 and CLOCK, which are central regulators of circadian rhythm, as well as SMAD3 and TGIF1, which mediate TGF-β signaling. This co-localization suggests that AHR may coordinate with multiple transcription factors at NC-XRE-rich regions, potentially forming regulatory complexes that extend beyond canonical AHR–ARNT interactions. Interestingly, 5x NC-XRE motifs exhibited stronger giggle overlap with AHR ChIP-seq datasets than 5x XRE motifs. We interpret this not as evidence that AHR globally prefers NC-XREs, but rather as a consequence of the relatively high frequency of clustered NC-XREs at known AHR binding sites in mouse liver. These findings reinforce the idea that NC-XRE clusters are enriched in regulatory DNA where AHR and its potential partners co-occupy, highlighting their functional importance.

Additionally, we observed significant overlap with several transcription factors identified through ORA and motif enrichment analyses in ChIP-seq datasets derived from non-liver mouse tissues (Supplementary Figure 4C), pointing to broader regulatory interactions that may extend beyond liver-specific contexts. These three complementary approaches—motif enrichment (Fig. 4A), regulon analysis (Fig. 4B), and ChIP-seq co-localization via giggle (Fig. 4C, 4D)—provide independent lines of evidence that converge on overlapping sets of transcription factors. This convergence increases confidence that the identified TFs are potential partners of AHR at NC-XRE loci, rather than artifacts of any single analysis.

We found several motifs enriched in peaks containing both XRE and NC-XRE motifs. As expected, the ARNT motif was enriched in both, supporting the previously known association with XRE motifs and suggesting that the AHR-ARNT complex could be associated with NC-XRE motifs (Fig 4A). However, ARNT was not enriched in our regulon ORA involving AHR peaks containing 5X NC-XRE motifs. This observation aligns with the findings of Huang and Elferink, who originally observed AHR bound to the NC-XRE motif in the promoter region of *Serpine1*, but did not detect ARNT (Huang and Elferink, 2012). This might indicate that AHR can function independently of ARNT at NC-XRE binding sites.

AHR is known to function through activation complexes involving various coregulators, and our findings suggest that several transcription factors may co-occupy AHR-bound genomic regions. Among these, BMAL1 (also known as ARNTL), a basic helix-loop-helix (bHLH) transcription factor like AHR, emerged as a particularly intriguing example. BMAL1 is well known for forming heterodimers with CLOCK to regulate circadian rhythm genes such as *Period* (*Per1, Per2, Per3*) and *Cryptochrome* (*Cry1, Cry2*) (Takumi, Matsubara, *et al*., 1998; Takumi, Taguchi, *et al*., 1998; Kume *et al*., 1999; Hogenesch *et al*., 1998). We observed co-occupancy of AHR ChIP-seq peaks containing 5X NC-XRE motifs with both BMAL1 and CLOCK binding sites (Fig 4D). We also saw that two BMAL1 target genes, *Noct* and *Serpine1*, are significantly overrepresented in the CollecTRI ORA analysis. These two target genes are also upregulated in mouse liver following TCDD exposure (Fig 3A, Fig 4B). Moreover, these AHR-5x NC-XRE peak regions also overlapped with ChIP-seq peaks for *Per2*, *Cry1*, and *Cry2*, suggesting that AHR may be involved in regulating components of the circadian clock. Our findings are consistent with a prior study demonstrating that activated AHR can interact with BMAL1 at E-box sequences and disrupt CLOCK-BMAL1–mediated transcription in cultured human hepatoma cells (Xu *et al*., 2010). Our data support and extend this work by showing genome-wide co-localization of AHR and BMAL1 binding sites in the mouse liver following TCDD exposure and suggest that AHR-BMAL1 interactions and subsequent signaling may be mediated through binding at NC-XRE motifs.

Several transcription factor motifs were specifically enriched in the regions flanking NC-XREs, but not in regions flanking canonical XREs. Among these, binding sites for TGIF1 and SMAD3 were particularly prominent (Figure 4A). Consistent with this, multiple targets of SMAD family transcription factors were upregulated following TCDD exposure in mouse liver (Figures 3A, 4A). Furthermore, both SMADs and TGIF1 non-liver ChIP-seq datasets showed significant overlap with AHR-NC-XRE peaks (Supplementary Figure 4C). These findings point to a potential crosstalk between AHR and other known signaling pathways, in this particular case, the TGF-β/SMAD signaling pathway at NC-XRE-enriched regulatory regions. Notably, such crosstalk has not been previously reported, highlighting its potential as a novel and meaningful observation. It suggests that AHR may integrate environmental signals with developmental and homeostatic pathways through non-canonical DNA binding mechanisms, expanding the scope of AHR’s regulatory network.

Other transcription factors such as KLF6 have been reported to associate with AHR at NC-XREs to regulate *Serpine1* expression (Wilson *et al*., 2013). Our findings are consistent with the idea that NC-XREs may provide a platform for diverse AHR-transcription factor interactions. For example, EP300, which is already known to interact with the AHR:ARNT complex (Kobayashi *et al*., 1997), showed strong peak overlap with AHR at 5x NC-XRE sites. We also saw overlap with other known co-regulators like E2F and HNF4A (Marlowe *et al*., 2004; Cholico *et al*., 2022). Although these proteins were already linked to AHR, their potential binding the at NC-XRE-rich regions has not been studied. Even more interesting, we found transcription factors, which have not been previously linked to AHR as a potential coregulator. These include factors involved in oncogenic signaling (e.g., MYC), and core transcriptional machinery (e.g., POLR2A, POLR2B). Their presence at the AHR-NC-XRE sites suggests that AHR may regulate cell cycle and RNA production using a previously uncharacterized non-canonical pathway.

### Testing NC-XRE function in a natural genomic context

Our bioinformatics analysis supports the hypothesis that AHR regulates target gene expression via NC-XRE DNA. To functionally test this hypothesis at a specific AHR target gene, we focused on *Serpine1*. ChIP studies in mouse liver identified an AHR binding site ∼150 bp upstream of the transcription start site, which contains a run of 5 NC-XRE motifs (Fig 5A) (Huang and Elferink, 2012). Indirect evidence suggests that AHR upregulates *Serpine1* expression via NC-XRE motifs, but this has never been tested directly.

**Figure 5.**
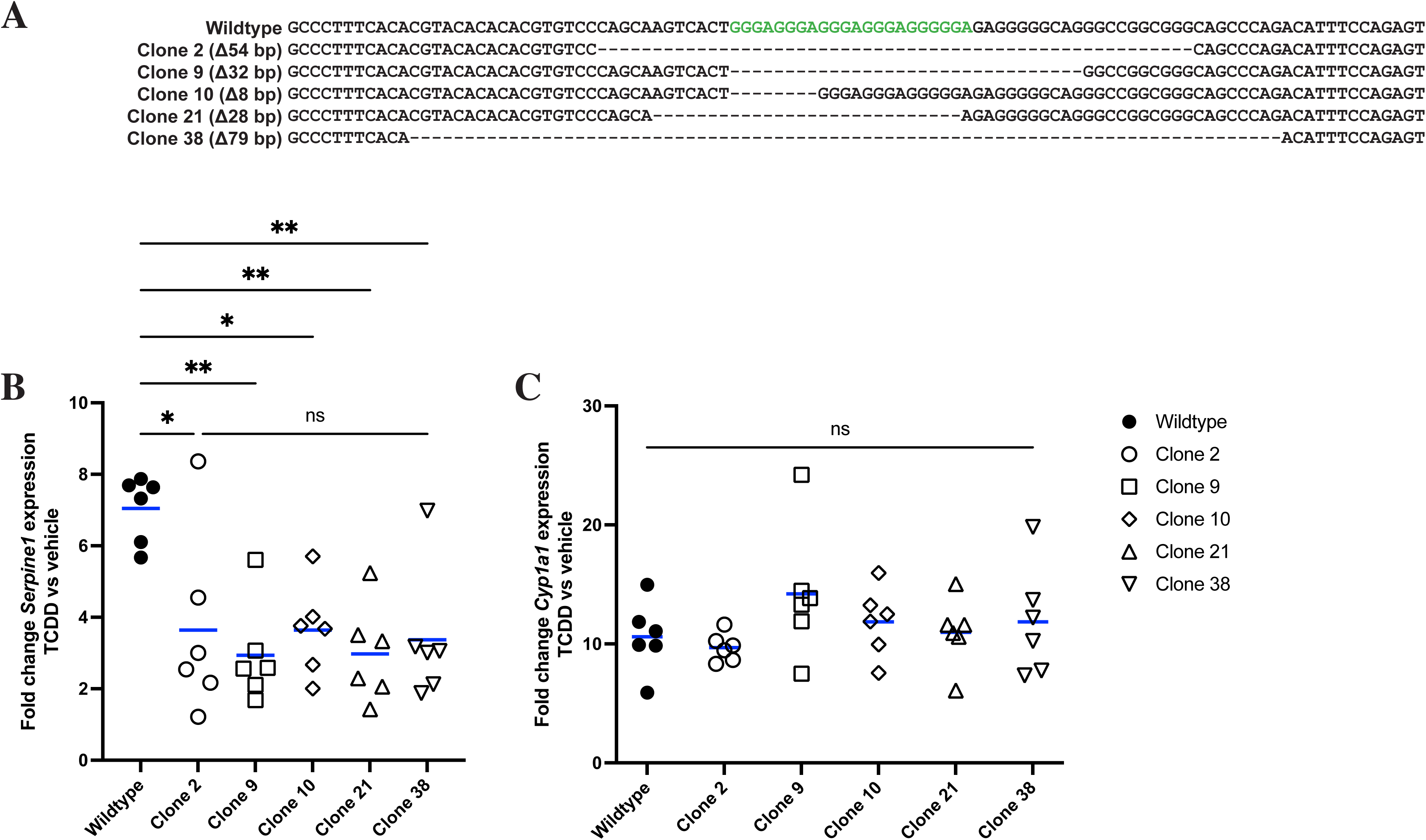
Mutations in NC-XRE DNA in *Serpine1* promoter reduced TCDD-dependent expression of *Serpine1*. (A) Schematic of mouse *Serpine1* gene showing AHR ChIP peak ∼150 bp upstream of transcription start site, which contains 5x repeated NC-XRE (nucleotides in red) and no XRE sequences. Hepa1-6 cell clones containing deletions in the NC-XRE region (dashed lines) are shown below the wild-type sequence. (B, C) Hepa1-6 wildtype or mutant cells were exposed to 10 nM TCDD or vehicle for 2 hours, followed by RNA extraction and qPCR for *Serpine1* (A) or *Cyp1a1* (B). Mutant cell lines with full or partial deletion of NC-XRE exhibit reduced increase in *Serpine1* following TCDD exposure compared to wild-type cells. Mutant cells showed similar increase in *Cyp1a1* following TCDD exposure as wild-type cells. Gene expression normalized to vehicle, 18S rRNA used as reference gene. One-way ANOVA with Tukey’s multiple comparisons test, * p < 0.05, ** p < 0.01, ns, not significant (p > 0.05).

We cultured mouse Hepa1-6 liver cells, used CRISPR-Cas9 to delete the 5x NC-XRE motif in the *Serpine1* promoter, generated clonal mutant cell lines, and directly tested whether the 5x NC-XRE motif contributes to TCDD-dependent *Serpine1* expression. To control for heterogeneity in gene expression between wildtype cells (Westermann *et al*., 2022; Giuliano *et al*., 2019), we first derived a population of wild-type Hepa1-6 cells from a single cell (see methods). These clonal wild-type cells were used for all subsequent experiments. We transfected Hepa1-6 cells with Cas9 protein together with a guide RNA targeting the NC-XRE sequence, picked single cells and established multiple cell lines containing different deletion mutations in the target NC-XRE (Fig 5A).

We exposed mutant and wild-type cells to TCDD or vehicle for 2 hours and assayed *Serpine1* expression by RT-qPCR. In all clones, deletion of the NC-XRE reduced upregulation of *Serpine1* in response to TCDD (Fig 5B). To test whether AHR function was normal, we also examined expression of *Cyp1a1*, a prototypical AHR target gene thought to be regulated by XRE motifs. All cell lines exhibited increased *Cyp1a1* expression following TCDD exposure, and there was no difference in mean *Cyp1a1* expression levels between wild-type and any mutant cell line (Fig 5C). Our results suggest that AHR transcription factor function is normal in NC-XRE mutant cell lines. We conclude that AHR upregulates *Serpine1* via this NC-XRE sequence 150 bp upstream of the transcription start site.

## DISCUSSION

We performed a genome-wide assessment of NC-XRE motifs and find they are prevalent in regulatory regions associated with AHR target genes. We also provide functional evidence that NC-XRE motifs in the *Serpine1* promoter are necessary for full upregulation of *Serpine1* in response to TCDD. Although deletion of all five NC-XRE motifs in the *Serpine1* promoter reduced TCDD-induced *Serpine1* expression by ∼50%, residual induction remained. This suggests that additional AHR-responsive elements, such as the two canonical XREs located ∼1000 bp upstream of the Serpine1 promoter, may also contribute to its regulation. Thus, NC-XREs and XREs likely act together to shape the overall transcriptional response of *Serpine1* to TCDD. However, we have only a basic understanding of the physiologic relevance of AHR signaling at NC-XRE motifs. Only two target genes (*Serpine1* and *Cdkn1a*) are known to be directly regulated by AHR at NC-XRE DNA, and these interactions were explored in only one cell type, hepatocytes, in adulthood (Huang and Elferink, 2012; Jackson *et al*., 2014). The degree to which other AHR target genes are regulated by non-consensus DNA sequence, in additional tissues and cell types and at different stages of organismal development, is not known.

To address this, we performed integrative pathway and transcription factor analyses of AHR target genes associated with 5x NC-XRE clusters. We identified 22 genes upregulated by TCDD that are proximal to these clusters, and pathway enrichment analyses revealed significant associations with biological processes such as development, morphogenesis, oxygen response, and enzymatic activity. These findings suggest that AHR may regulate a broader array of physiological processes through NC-XRE-mediated transcriptional control than previously appreciated.

ARNT is considered an obligate binding partner of AHR (Reyes *et al*., 1992), but these studies focused on AHR-ARNT interactions at XRE DNA. Whether ARNT binds AHR at NC-XRE DNA and whether NC-XRE– mediated transcriptional responses require ARNT has not been determined. To our knowledge, no studies have directly tested NC-XRE–dependent gene regulation in ARNT-deficient cell lines, and this remains an important question for future research. Studies suggest that under certain conditions AHR can act independently of ARNT to upregulate gene expression. In mouse liver, AHR was bound to NC-XRE in the promoter of the *Serpine1* gene following TCDD exposure, but no ARNT was detected (Huang and Elferink, 2012). In a rat hepatoma cell line containing a Serpine1:luciferase reporter construct, TCDD increased luciferase levels but knockdown of ARNT had no effect (Huang and Elferink, 2012). Our analysis supports this observation. Although ARNT motifs were enriched in AHR peaks containing both XRE and NC-XRE motifs, ARNT did not show co-occupancy at AHR peaks containing 5x NC-XRE motifs. This raises the possibility that AHR may function independently of ARNT at NC-XRE sites, potentially through interactions with alternative co-regulators.

A limitation of the previous studies is that none considered ARNT2. ARNT2 shares significant sequence homology with ARNT and was shown to interact with AHR and XRE DNA *in vitro* (Hirose *et al*., 1996; Tanguay *et al*., 2000; Rowatt *et al*., 2003), but to our knowledge there is absence of evidence whether ARNT2 interacts with AHR *in vivo*. DNA binding, coregulator recruitment and target gene expression could be different between AHR-ARNT and AHR-ARNT2 transcriptional complexes. Given our findings that ARNT1 does not appear to co-occupy AHR-bound NC-XRE regions, it is plausible that ARNT2 may serve as an alternative co-regulator in these contexts. Notably, we were unable to assess ARNT2 co-localization at AHR-NC-XRE peaks due to the absence of publicly available ARNT2 ChIP-seq datasets. As such, the potential involvement of ARNT2 in AHR-NC-XRE-mediated transcription remains an open question that warrants further study. This raises the intriguing possibility that AHR may form distinct transcriptional complexes depending on the DNA motif it binds—partnering with ARNT1 at XREs and potentially with ARNT2 at NC-XREs. Such differential dimerization could influence target gene specificity, chromatin recruitment, and transcriptional outcomes, warranting deeper investigation into the role of ARNT2 in non-canonical AHR signaling. However, expression of Arnt2 in mice is largely restricted to neuronal tissues and the developing kidney, with little to no detectable expression in mouse liver (Keith *et al*., 2001; Aitola and Pelto-Huikko, 2003). Consistent with this expression pattern, overexpression of Arnt2 in hepatic cells produces minimal TCDD responsiveness compared to Arnt (Dougherty and Pollenz, 2008). Together, this suggests that while Arnt2 could influence AHR-NC-XRE-mediated transcription, it is unlikely to function as an AHR binding partner in the liver.

Our integrative analysis uncovered a broader network of transcriptional regulators co-localizing at AHR-NC-XRE sites, including factors involved in circadian rhythm (e.g., Arntl/Bmal1, Clock), TGF-β/SMAD signaling (e.g., SMAD3, TGIF1), and oncogenic or epigenetic regulation (e.g., MYC, EP300, BCL6). These findings suggest that AHR may orchestrate complex transcriptional programs through non-canonical DNA binding, extending its influence into diverse biological pathways.

## LIMITATIONS

The evidence for transcription factor co-occupancy at AHR-bound NC-XRE clusters is indirect. We compared AHR peaks from TCDD-treated liver with transcription factor peaks obtained from untreated liver or hepatocyte datasets, which do not provide temporal or spatial resolution of binding events. It, therefore, remains possible that nearby transcription factors occupy adjacent loci without functionally interacting with AHR. For example, a canonical E-box motif (CACGTG, recognized by CLOCK-BMAL1 and other bHLH factors) lies immediately upstream of the NC-XRE cluster in the *Serpine1* promoter. Yet deletion of this sequence in clone 38 had no additional effect on TCDD-induced *Serpine1* expression beyond NC-XRE deletion alone. This indicates that, at least for *Serpine1*, the E-box is dispensable for AHR-dependent regulation and that co-occupancy with CLOCK-BMAL1 is unlikely to explain the observed transcriptional response. These findings underscore that functional validation at individual loci is essential. Direct demonstration of simultaneous AHR and co-factor binding will require future experiments such as sequential ChIP, proximity ligation assays, or single-molecule imaging.

The computational analyses performed here suggest many hypotheses, such as that AHR target genes may be regulated by NC-XRE DNA. However, the algorithms employed here make predictions but lack experimental validation. We demonstrated that the repeated NC-XRE motifs in the promoter region of *Serpine1* regulate AHR-dependent *Serpine1* expression in a mouse liver cell line. A recent study in human lung cancer cells showed upregulation of SERPINE1 by AHR agonists, suggesting SERPINE1 is also an AHR target in humans (Gorelik *et al*., 2025). Nevertheless, it remains unclear whether AHR regulates gene expression in humans via NC-XRE DNA motifs, as the human *SERPINE1* promoter contains only three GGGA repeats separated by 50 bp spacers. However, we appreciate that DNA regulatory elements may be incompletely conserved between species. Showing that AHR binds NC-XRE DNA to regulate gene expression in any species is important foundational knowledge. While the specific number or configuration of NC-XRE sequence at a given promoter or enhancer may differ between strains of animals or between species, it is extremely unlikely that AHR regulates gene expression by binding to NC-XRE in mice but not in humans. Considering that AHR binds identical XRE DNA sequence in zebrafish, mice and humans (Tanguay *et al*., 1999; Andreasen *et al*., 2002; Denison *et al*., 1988; Yao and Denison, 1992), we argue that the same is likely true for NC-XRE.

## METHODS

### Re-analysis of published murine liver ChIP-Seq datasets

ChIP-Seq datasets of AHR in murine livers after 2hrs of TCDD treatment in male (GSE97634) and female (GSE97636) mice were downloaded from NCBI Gene Expression Omnibus (GEO) (Table 1) (Fader *et al*., 2017, 2019; Nault *et al*., 2016). Data were trimmed using trim_galore (Krueger, 2015), mapped to the mouse genome build UCSC mm10 using bowtie2 (Langmead and Salzberg, 2012), and peaks were called using MACS2 using the provided controls (Langmead and Salzberg, 2012; Zhang *et al*., 2008). Peaks were merged using bedtoools into male, female and male + female merged datasets (Quinlan and Hall, 2010). Each peak was searched for AHR motifs using an in-house script. To test the likelihood of finding AHR motifs anywhere in the genome, random DNA peaks were generated matching the chromosome and size distribution as previously described (Coarfa *et al*., 2020), then searched for AHR motifs using a custom Ruby code. We analyzed AHR ChIP-seq peaks by extracting the underlying genomic sequences and scanning them for both XRE and NC-XRE motifs. Motifs provided in IUPAC notation were converted into regular expressions, allowing systematic identification and annotation of motif occurrences within peak regions (doi.org/10.5281/zenodo.17211132). For each motif of interest (XRE, NC-XRE, RelBAhRE), we compared the total number of motif-containing peaks in the observed dataset against the matched random set using a Fisher’s exact test. Since only three tests were performed, we did not perform multiple testing correction and reported raw Fisher’s exact test p-values with significance defined at p < 0.05. Venn diagrams of AHR motifs in the merged AHR peaks were derived using *eulerr* package (J, 2024) in R studio (RStudio Team, 2025) and pie charts were made using using GraphPad Prism version 10.2.2, MacOS, GraphPad Software, Boston, Massachusetts, USA, www.graphpad.com.

To investigate the genomic distribution of AHR binding sites, we used ChromHMM, a chromatin state annotation tool (Ernst and Kellis, 2012). We utilized ChIP-Seq datasets for eight histone modifications— H3K27ac, H3K27me3, H3K36me3, H3K79me2, H3K9ac, H3K4me3, H3K4me1, and H3K9me3—along with an input control, all derived from 8-week-old mouse liver tissue. These datasets were downloaded from the NCBI Gene Expression Omnibus (GEO; accession numbers: GSM1000140, GSM918718, GSM1000150, GSM1000151, GSM1000152, GSM1000153, GSM769014, GSM769015, GSM751035) (Table 1) (Fader *et al*., 2017, 2019; Nault *et al*., 2016). Raw reads were trimmed using *trim_galore* (Krueger, 2015) and aligned to the UCSC mm10 mouse genome build using *bowtie2* (Langmead and Salzberg, 2012), The aligned reads were converted into binned signal representations alongside genomic windows of signal abundance, which were then used to train a hidden Markov model to segment the genome into 10 chromatin states (Ernst and Kellis, 2017) using the ChromHMM tool (Ernst and Kellis, 2012). We overlaid these chromatin states onto AHR peaks containing XRE and NC-XRE motifs, as shown in Figures 1C, 1D, and 2D, using a custom in-house script. Additionally, we used the same histone ChIP-Seq datasets to compute signal distributions across genomic regions (Figure 2E) using the HOMER software (Heinz *et al*., 2010). All figures were generated using GraphPad Prism version 10.2.2.

Runs of AHR motifs, e.g. sequences of successive motifs within a maximum distance from each other, were determined using an in-house Python script and bedtoools. Flanking sequences were determined using the bedtools software (Quinlan and Hall, 2010). Enriched motifs within the flanking sequences were determined using the HOMER software (Heinz *et al*., 2010).

To identify transcription factors with potential co-binding at AHR-bound NC-XRE or XRE clusters, we performed giggle score analysis using the CistromeDB Toolkit (Zheng *et al*., 2019; Mei *et al*., 2017). As input, we used mouse liver AHR ChIP-seq peaks (GSE97634, GSE97636) containing either 5 canonical XRE motifs or 5 NC-XRE motifs with a maximum spacer of 50 base pairs. Giggle calculates an integrated score that combines statistical significance (Fisher’s exact test p-value) and enrichment (odds ratio) to quantify whether two ChIP-seq peak sets overlap more than expected by chance (Layer *et al*., 2018).

The giggle score thus reflects both the strength and significance of overlap: higher scores indicate stronger co-localization between two datasets, consistent with the possibility that the corresponding proteins bind the same genomic loci and may participate in transcriptional complexes.

Although giggle automatically compares query peaks against all publicly available datasets in the Cistrome Data Browser, for interpretation we filtered the results to include only ChIP-seq datasets derived from mouse liver tissue or hepatocyte-derived cell lines. This restriction ensured that the comparisons were performed in a biologically relevant context.

The giggle scores were visualized in Figure 4C–D to illustrate overlap with AHR datasets (serving as an internal control) and with additional transcription factors that may act as potential AHR co-regulators.

### Re-analysis of published murine liver AHR RNA-Seq datasets and integration with ChIP-seq datasets

Several RNA-seq datasets of mouse liver after treatment with TCDD at different doses and for multiple timepoints were downloaded from NCBI GEO: GSE109863, GSE62902, and GSE87519 (Nault *et al*., 2018, 2015; Fader *et al*., 2017) (Table 1). Data was trimmed using trim_galore (Krueger, 2015), then mapped onto the mouse genome build UCSC mm10 using STAR (Dobin *et al*., 2013), and gene expression was quantified using feature_counts (Liao *et al*., 2014). Differentially expressed genes between TCDD and vehicle treatment were inferred using the EdgeR and RUVr R packages (Robinson *et al*., 2010; Risso *et al*., 2014), with significance achieved at fold change exceeding 1.5x and FDR-adjusted p-value<0.05. We used the ranked gene list to perform Gene Set Enrichment Analysis (Subramanian *et al*., 2005) using datasets from mouse molecular signature database version 7.5.1 (Castanza *et al*., 2023). Genomic locations (represented as bed files) and gene signatures were integrated using bedtools (Quinlan and Hall, 2010). To capture both promoter and proximal enhancer effects, peaks were considered as associated with genes if the gene body was within 10,000 basepairs of a peak. Over Representation Analysis (ORA) was performed using a custom Python script, which applies the hypergeometric test used by MSigDB to assess enrichment of gene sets from the gene ontology biological process gene set version 7.5.1 (Castanza *et al*., 2023) and CollecTRI version 2 regulons (Müller-Dott *et al*., 2023).

### Cell culture

The mouse hepatoma cell line Hepa 1–6 was obtained from ATCC (Manassas, VA) (Darlington *et al*., 1980). Hepa 1-6 cells cells were cultured in 10 cm plates in Dulbecco’s Modified Eagle’s medium (DMEM) supplemented with 1% (v/v) Penicillin/Streptomycin and 10% (v/v) FBS in a humidified incubator (5% CO2, 37°C). To derive wild-type clones for CRISPR mutagenesis, Hepa 1-6 cells were separated using BD FACSAria II cell sorter (Cytometry and Cell Sorting Core, Baylor College of Medicine) and one cell per well was plated into a 96-well plate. Cells grew for 2-3 weeks until colonies were visible and selected for further expansion. Wild-type clone 12 was used for subsequent studies.

### CRISPR mutagenesis of Hepa 1-6 cells

We used CHOPCHOP (Labun *et al*., 2019) to design guide RNAs targeting the 5x NC-XRE motif region 150bp upstream of transcription start in the mouse Serpine1 gene (target sequence CAGCAAGTCACTGGGAGGGA<u>GGG</u>, PAM motif underlined). Synthetic single guide RNA—modified with 2’-O-methyl at 3 first and last bases, and 3’-phosphorothioate bonds between first 3 and last 2 bases—and purified SpCas9-2NLS protein was purchased from Synthego (Redwood City, CA). We mixed guide RNA and Cas9 (1.3:1 ratio) to form ribonucleoproteins according to Synthego’s recommended protocol. Guide RNA and Cas9 ribonucleoprotein complexes in serum-free medium were transfected into Hepa1-6 WT clone 12 cells (see above) using Lipofectamine^TM^ CRISPRMAX^TM^ Cas9 Transfection reagent (Thermo Fisher Scientific, Waltham, MA) according to the manufacturer’s protocol. 100,000 Hepa-1-6 WT-12 cells in standard cell culture media (above, containing FBS and antibiotics) were mixed with the ribonucleoprotein-transfection solution and split into two wells for genomic analysis and clonal expansion. Cells were incubated in a humidified incubator (5% CO2, 37 °C) for 24 hours. We then changed the media and allowed the cells to incubate for another 3 days before genotyping or deriving single-cell colonies for clonal expansion.

To derive mutant colonies from single cells, we separated and plated transfected WT-12 cells using BD FACSAria II cell sorter as above. 48 single cell clones were expanded and genotyped, focusing on mutant cell lines from clones 2, 9, 10, 21, and 38.

### Genotyping cell lines

Genomic DNA was extracted from cells using phenol-chloroform extraction. Cells were lysed with DNA lysis buffer (10 mM Tris, 200 mM NaCl, 5 mM EDTA, 1% SDS, 0.4 mg/ml proteinase K). Equal volume of phenol:choloroform:isoamyl alcohol was added for DNA extraction into upper phase followed by ethanol-sodium acetate precipitation of DNA. We used PCR to amplify the region flanking the NC-XRE motifs using primers 5’- AAGCCAGGCCAACTTTTCCT and 5’- CGGTCCTCCTTCACAAAGCT. PCR products were Sanger sequenced to identify mutations. To identify mutations from mosaic cell populations, we used ICE CRISPR Analysis Tool (Conant *et al*., 2022) to deconvolute Sanger sequencing results.

Mutant clones (all derived from WT clone 12), selected for further experiments, were clone 2 (54 bp deletion in NC-XRE region of Serpine1 promoter), clone 9 (32 bp deletion), clone 10 (8 bp deletion), clone 21 (28 bp deletion) and clone 38 (74 bp deletion). The deletions in each clone were further validated by next generation sequencing. We used PCR to amplify a 450 bp region of DNA encompassing the NC-XRE. After PCR amplification, DNA was quantified by Qubit dsDNA HS fluorometric quantitation (ThermoFisher Q32851). Cytation 5 imaging reader (BioTek; Santa Clara, CA) was used to read the samples. 500 ng of each amplicon was sent for Amplicon E-Z Next Generation Sequencing (GENEWIZ from Azenta Life Sciences, South Plainfield, NJ), paired end Illumina MiSeq 2X250 bp, 50,000 reads per sample. Each clone contained > 99% of the expected mutation.

### Quantitative reverse-transcription PCR

70-80 % confluent Hepa-1-6 parental WT12 cells and WT12 mutant clonal cell lines 2, 9, 10, 21, 38 were treated with 10 nM TCDD (n=6 per treatment group) or 0.1 % DMSO (vehicle control, n=6 per treatment group) and for 2 hours each. We selected 10 nM TCDD because this concentration has been shown to robustly activate AHR target genes while avoiding overt cytotoxicity (Humphrey-Johnson *et al*., 2015). Cells were then lysed in TRIzol followed by RNA extraction according to the manufacturer’s protocol using Direct-zol RNA Miniprep Kit, including the optional on-column DNase digestion (cat no, 11-331; Zymo Research, Irvine, CA).

RNA concentration was measured using NanoDrop 2000 Spectrophotometer (Thermo Scientific, Waltham, MA). Purity of the RNA (A260/A280) was ≥ 2 with the yield of 60ng/μl or higher. 1000 ng of the RNA were converted into cDNA using iScript Reverse Transcription Supermix for RT-qPCR (Biorad, Hercules, CA; cat no 1708841) in a total 20 μl volume and incubated for 5 min at 25°C followed by incubation at 46°C for 20 min, at 95°C for 1 min in C1000 Touch Thermal Cycler (Biorad, Hercules, CA).

To validate differential gene expression, we used real-time quantitative PCR (qPCR) by the PrimePCR Assay (Biorad, Hercules, CA) based PCR amplification using the CFX Opus 96 Real-Time PCR System (Biorad, Hercules, CA) via the 2^−ΔΔCt^ method under default reaction conditions (95°C for 2min; 95°C for 5 sec; 60°C for 30 sec. for 40 cycles). An Aliquot of 4 μl of 1:10 dilution of cDNA was mixed with 10 μL BIORAD SsoAdvanced Universal Probes Supermix (cat no 1725280) and target gene specific PrimePCR Assays (1 μL) and Nuclease free water up to a total volume of 20 μl (three technical replicates were run for each sample). cDNA samples were also used for measurement of Gapdh levels for normalization. Ct values were calculated by the Bio-Rad instrument using the CFX Maestro™ Software version 2.3.

The following PrimePCR Assays were used. Note that BioRad does not provide specific primer/probe sequences. BioRad provides the amplicon sequence with additional base pairs added to the beginning and/or end of the sequence (amplicon context sequence). This is in accordance with the minimum information for the publication of real-time quantitative PCR experiments (MIQE) guidelines (Bustin *et al*., 2011).

Serpine1, Mouse (qMmuCIP0037358)

**<u>Amplicon Context Sequence:</u>**

CTGTCGGGTTGTGCCGAACCACAAAGAGAAAGGATCGGTCTATAACCATCTCCG TGGGGGCCATGCGGGCTGAGATGACAAAGGCTGTGGAGGAAGACGCCACTGTG CCGCTCTCGTTTACCTCGATCCTGACCTTTTGCAGTGCCTGTGCT

**<u>*Gapdh*, Mouse (qMmuCEP0039581)</u>**

TGGGAGTTGCTGTTGAAGTCGCAGGAGACAACCTGGTCCTCAGTGTAGCCCAAGATGCCCTTCAGTGGGCCCTCAGATGCCTGCTTCACCACCTTCTTGATGTCA

***<u>Cyp1a1</u>*<u>, Mouse (qMmuCEP0041980)</u>**

CAGCCACCTAGATCATGCCTTCCATGTATGGACTTCCAGCCTTCGTGTCAGCCA CAGAGCTGCTCCTGGCTGTCACCGTATTCTGCCTTGGATTCTGGGTGGTCAGAG CCACAAGAACCTGGGTTCCCAAAGGCCTGAAGACTCC

The results shown were the average of three or more biologically independent experiments. The cumulative results were statistically analyzed by using Prism version 10 (GraphPad Software, Boston, MA).

## ACKNOWLEDGMENTS

We dedicate this manuscript to our dear friend and colleague Professor Cornelis Elferink, who provided invaluable advice and encouragement as we pursued these studies. Professor Elferink, who discovered the association between AHR and NC-XRE DNA, died unexpectedly during the preparation of this manuscript. TDP, SLG and CC were partially supported by The Cancer Prevention Institute of Texas (CPRIT) [RP210227, RP200504], NIH/NCI P30 shared resource grant [CA125123], NIH/NIEHS center grants [P30 ES030285] and [P42 ES027725], and NIH/NIMHD [P50MD015496]. DAG supported by NIEHS R01 ES026337.

---

**Supplementary Figure 1.**
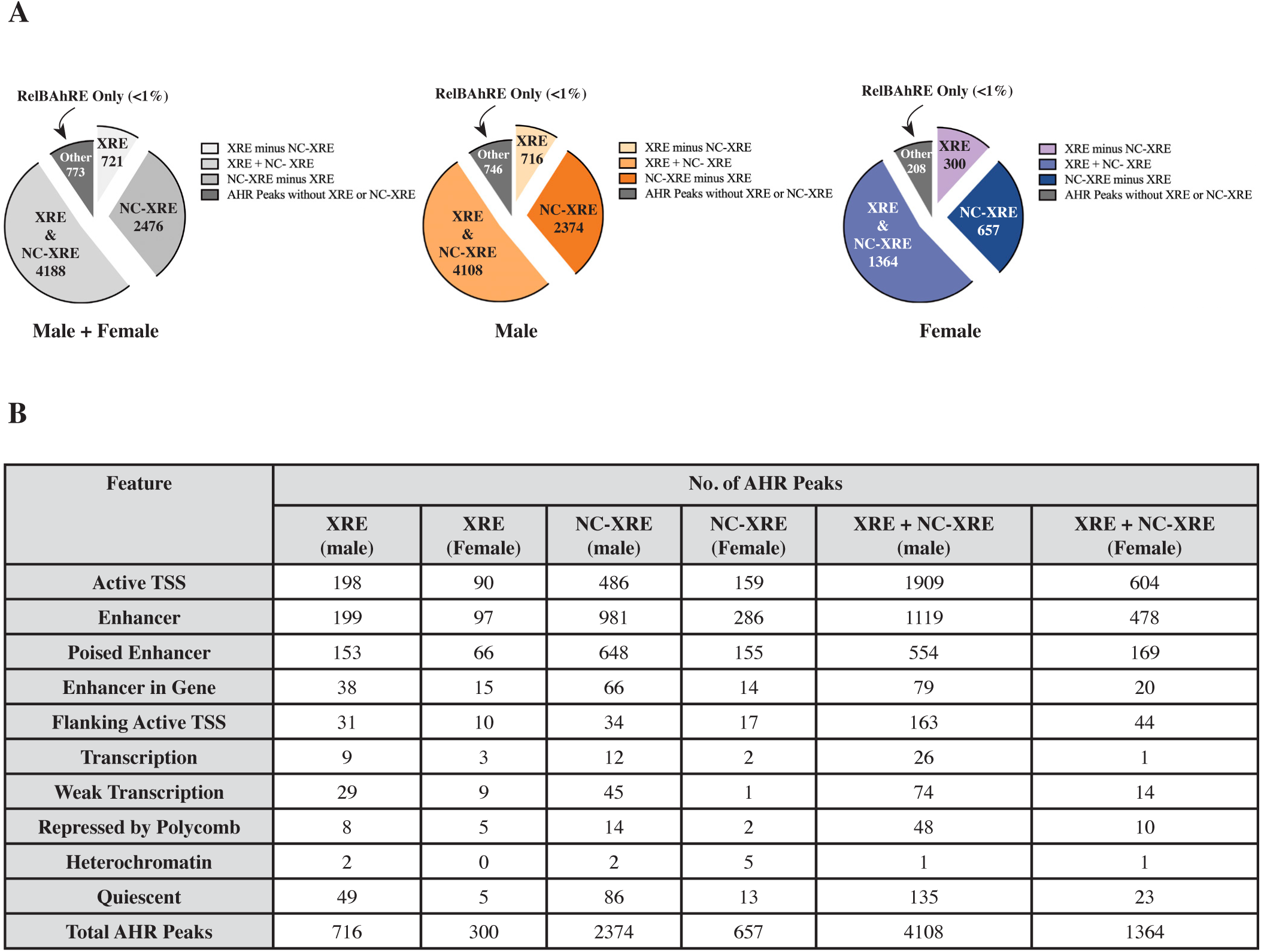
Distribution of AHR ChIP-seq Peaks and Chromatin States. **(A)** Pie chart showing the distribution of AHR ChIP-seq peaks from mouse liver containing XRE, NC-XRE, and RelBAhRE motifs, corresponding to data in Figure 1B. The chart highlights that the proportion of XRE and NC-XRE motifs are similar between the sexes. The number of RelBAhRE motifs is miniscule compared to XRE and NC-XRE motifs. **(B)** Table detailing the genomic distribution of AHR ChIP-seq peaks across various chromatin states, corresponding to Figure 1***(C-D)***. It includes the number of AHR peaks associated with each chromatin state for different motif variations (XRE/ NC-XRE for male/ female/ combined datasets). Chromatin states include Active TSS, Enhancer, Poised Enhancer, Weak Transcription, Repressed by Polycomb, Heterochromatin, and Quiescent regions.

**Supplementary Figure 2.**
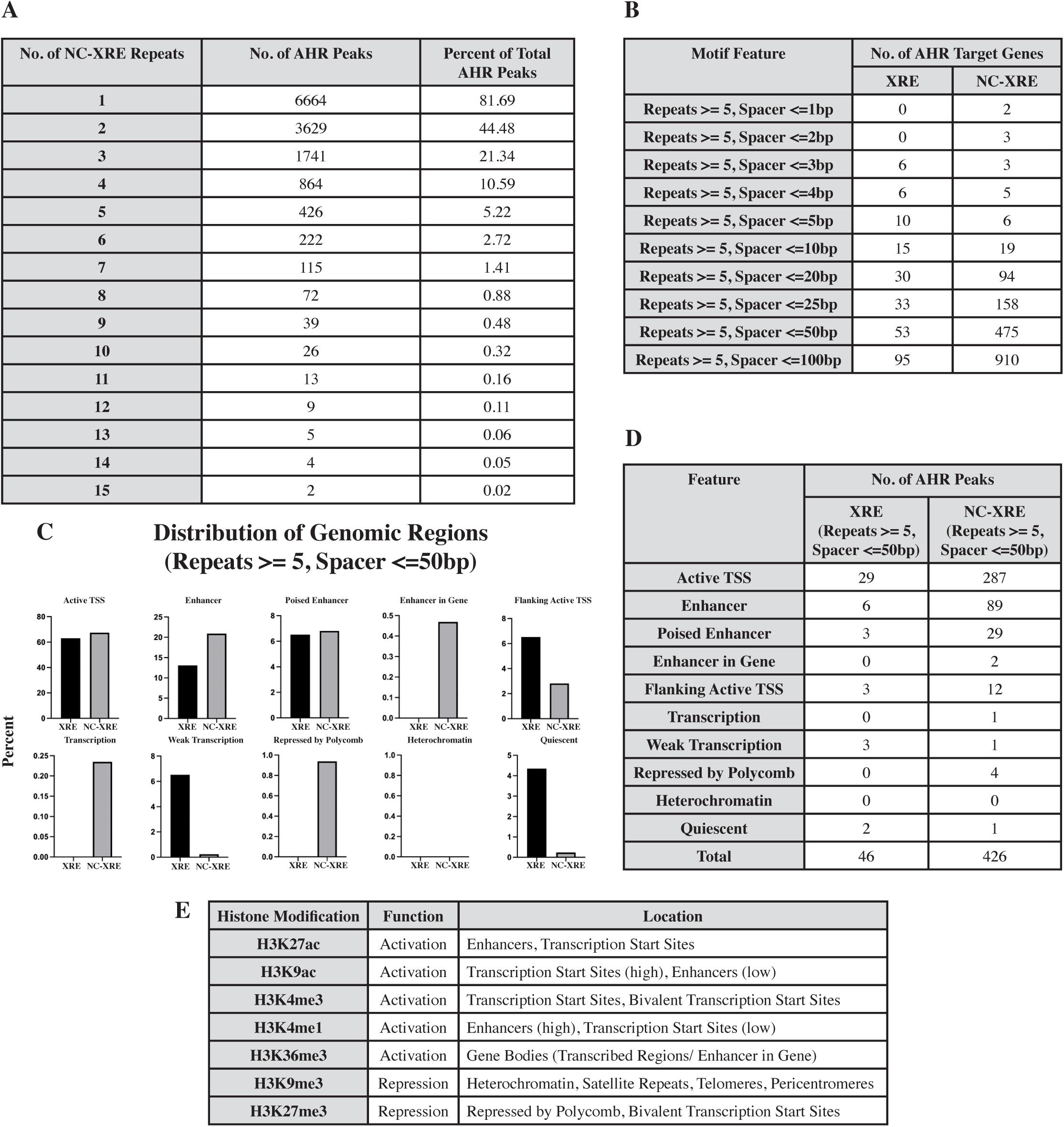
Detailed analysis of NC-XRE repeats and their genomic context. (A) Number of AHR ChIP peaks containing NC-XRE repeats (1-15) separated by 50 bp or less, corresponds to Figure 2C. (B) Number of putative AHR target genes within 10 kb of AHR ChIP-seq peaks containing 5 repeats of XRE/NC-XRE motifs separated by different numbers of spacers, corresponds to Figure 2B. (C) Enlarged view of Figure 2D, for detailed comparison of chromatin state distributions. 5 repeats of XRE or NC-XRE motifs with 50 bp spacers are mostly present near transcription start sites and enhancers. (D) Values for the data shown in (C). (E) Overview of histone modifications, indicating their functions (activation or repression) and typical locations (e.g., transcription start sites).

**Supplementary Figure 3.**
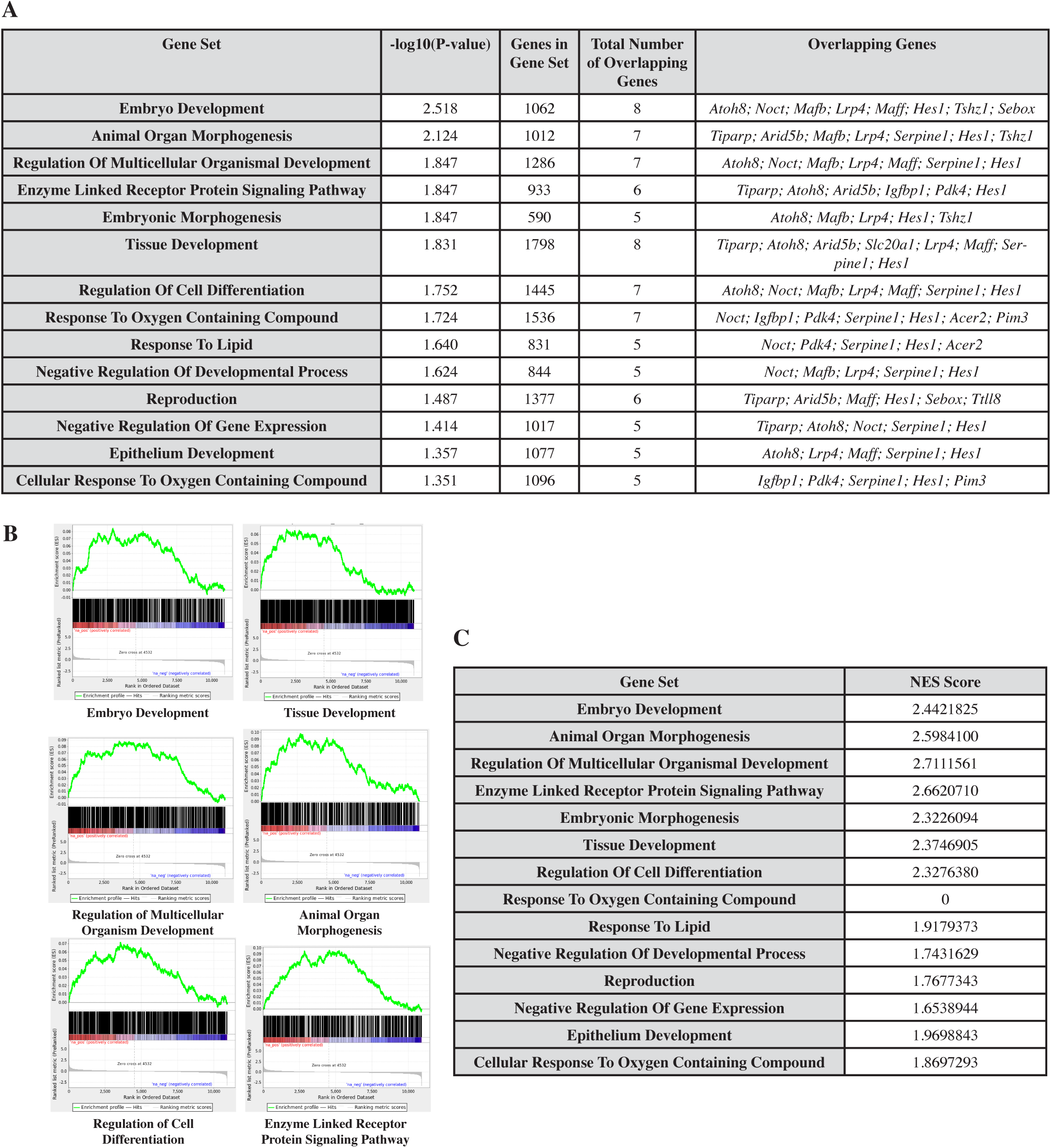
Enrichment analysis of putative AHR-NC-XRE target genes. **(A)** Table corresponding to the ORA data shown in Figure 3B, including the enriched Gene Ontology Biological Processes (GOBP) gene sets and overlapping genes that have at least 5 NC-XRE repeats with a maximum of 50 bp spacers within 10 kb of the gene body. **(B)** GSEA enrichment plots of 6 upregulated pathways identified in the ORA analysis. These panels are ranked based on the number of hit genes in the ORA: Embryo development (8), tissue development (8), regulation of multicellular organismal development (7), animal organ morphogenesis (7), regulation of cell differentiation (7), and enzyme-linked receptor protein signaling pathway (6). **(C)** Normalized enrichment scores (NES) of the GOBP gene sets shown in Figure 3D.

**Supplementary Figure 4.**
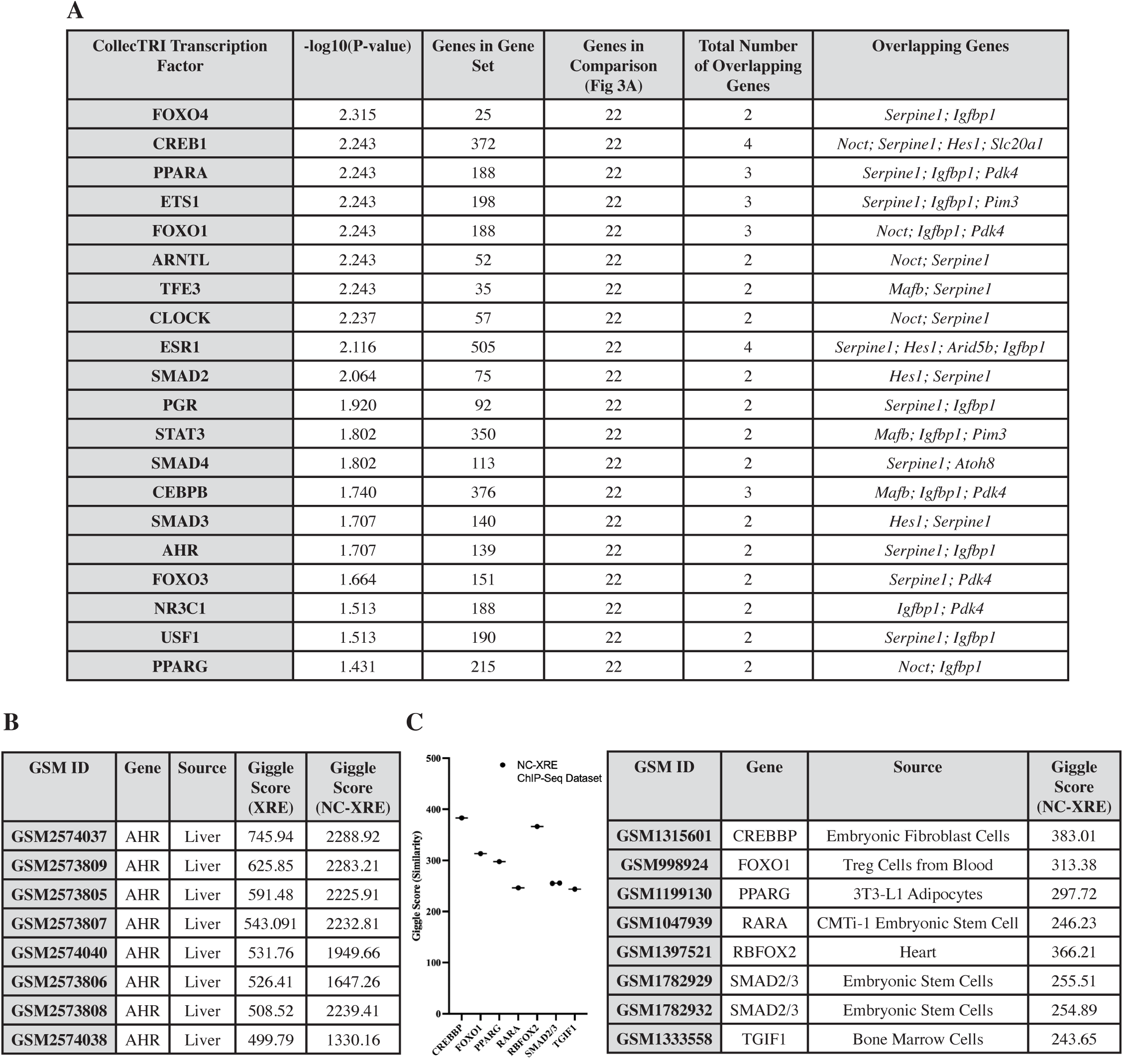
Putative transcription factors targeting 5 repeats of XRE or NC-XRE separated by 50 bp or less. (A) Corresponds to the ORA data shown in Figure 4B, including the upregulated CollecTRI transcription factors and overlapping genes that have 5 NC-XRE repeats with a maximum of 50 bp spacers within 10 kb of the gene body. (B) Corresponds to the CistromeDB giggle scores shown in Figure 4C, indicating the overlap of AHR peaks containing 5 or more XRE or NC-XRE with 50 bp spacers in our query dataset compared mouse liver AHR ChIP-seq datasets on the CistromeDB database. Higher giggle score means higher similarity. (C) Giggle scores of AHR peaks with 5 or more NC-XRE repeats with up to 50 bp spacers compared to some of the hit transcription factor (Fig 4A, 4B) peaks from mouse non-liver ChIP-seq experiments. Each point represents how strongly NC-XRE containing AHR binding sites overlap with the transcription factor binding sites. A higher giggle score indicates a higher likelihood of these transcription factors co-occupying the AHR peaks.

**Supplementary Figure 5.**
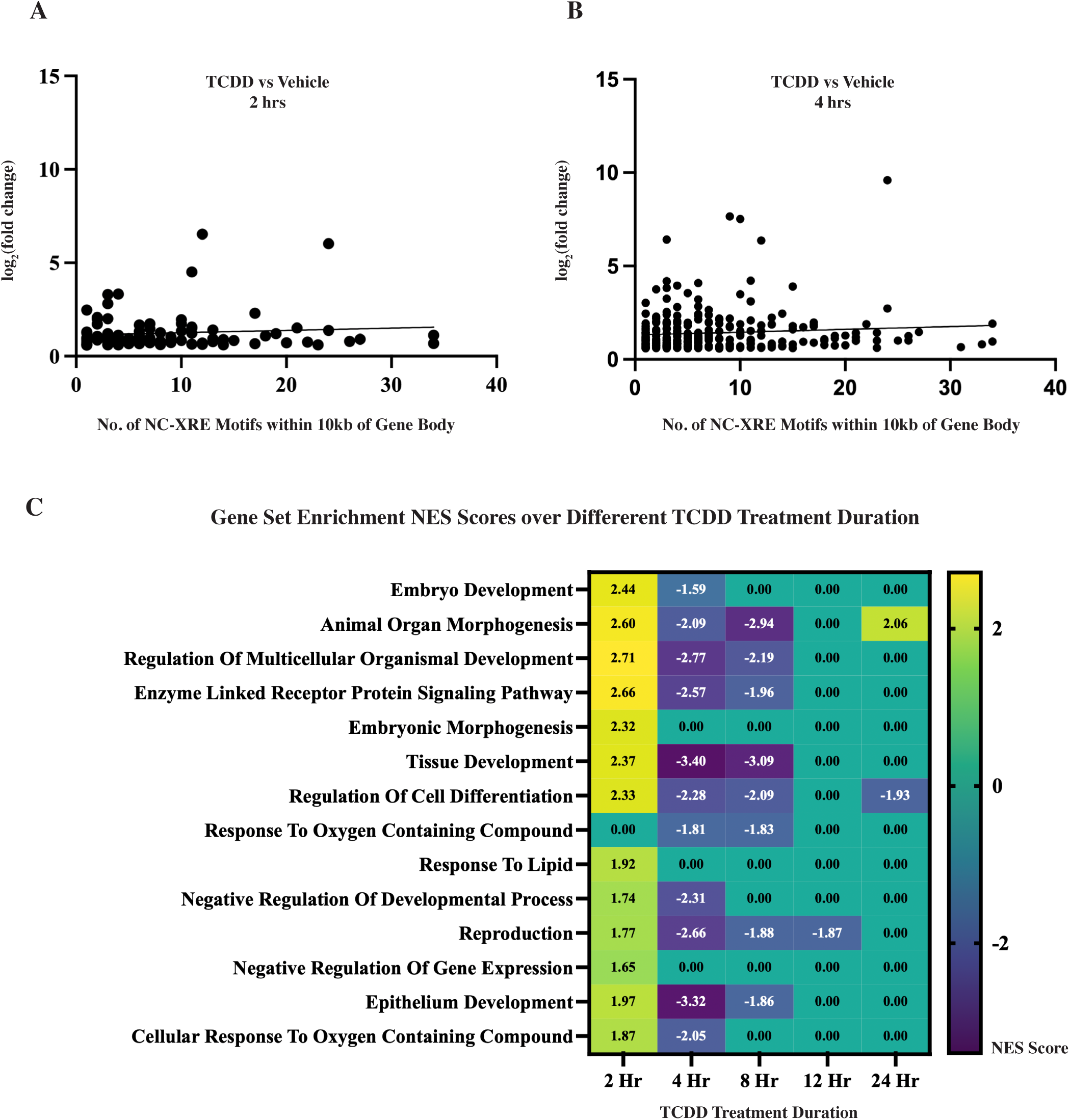
Relationship between NC-XRE motif number, TCDD treatment duration, and AHR target gene expression. (A,B) Simple linear regression of upregulated AHR target genes containing at least one NC-XRE motif within 10 kb of the gene body after 2h (A) and 4h (B) of TCDD treatment. Each point represents a single gene. The x-axis indicates the number of NC-XRE motifs, and the y-axis shows log2(fold change) of genes (TCDD vs. vehicle). No significant linear relationship was observed between motif number and gene expression (2 h: p = 0.3653; 4 h: p = 0.1430) at either timepoint. (C) Gene Set Enrichment Analysis (GSEA) of global mouse liver RNA-seq (TCDD vs. DMSO, ranked gene lists) across multiple treatment durations. Only pathways identified in the ORA (Fig. 3B) are shown. At 2h, 13 of 14 ORA-identified pathways were significantly upregulated (NES, FDR < 25%), followed by a shift toward indirect or secondary AHR target pathways with longer treatment durations, resulting in a predominant downregulation of these pathways.

**Supplementary Table 1: AHR Target Genes with Different Combinations of XRE or NC-XRE Near Gene Body.** Frequency of repeated runs of XRE or NC-XRE motifs within 10000 bp of a gene body. It shows the total number of AHR ChIP peaks containing 5 repeats of XRE or NC-XRE separated by spacers ranging from 1 to 100 basepairs (bp) along with the expression pattern (log2FoldChange) of a particular gene (Fold Change > 1.5x; FDR <0.05).

**Supplementary Table 2. Similarity of AHR ChIP-Seq Peaks containing 5 NC-XRE with Known Transcription Factor Peaks.** Giggle scores of AHR peaks with 5 or more NC-XRE repeats with up to 50 bp spacers compared to known transcription factor peaks from mouse liver or liver cell ChIP-seq datasets. A higher giggle score indicates a higher likelihood of these transcription factors co-occupying the AHR peaks.

## Notes

### Competing Interest Statement

DAG is the editor in chief of the journal Biology Open, published by the nonprofit The Company of Biologists.

### Summary of Updates

Clarifications made to the methods, added supplemental figure 5

## REFERENCES

Aitola, M.H. and Pelto-Huikko, M.T. (2003) Expression of Arnt and Arnt2 mRNA in developing murine tissues. J. Histochem. Cytochem., 51, 41–54.

Andreasen, E.A. et al. (2002) The zebrafish (Danio rerio) aryl hydrocarbon receptor type 1 is a novel vertebrate receptor. Mol. Pharmacol., 62, 234–249.

Beischlag, T.V. and Perdew, G.H. (2005) ER alpha-AHR-ARNT protein-protein interactions mediate estradiol-dependent transrepression of dioxin-inducible gene transcription. J. Biol. Chem., 280, 21607–21611.

Bustin, S.A. et al. (2011) Primer sequence disclosure: a clarification of the MIQE guidelines. Clin. Chem., 57, 919–921.

Castanza, A.S. et al. (2023) Extending support for mouse data in the Molecular Signatures Database (MSigDB). Nat. Methods, 20, 1619–1620.

Cholico, G.N. et al. (2022) Genome-Wide ChIPseq Analysis of AhR, COUP-TF, and HNF4 Enrichment in TCDD-Treated Mouse Liver. Int. J. Mol. Sci., 23.

Coarfa, C. et al. (2020) Epigenetic response to hyperoxia in the neonatal lung is sexually dimorphic. Redox Biol., 37, 101718.

Conant, D. et al. (2022) Inference of CRISPR Edits from Sanger Trace Data. The CRISPR Journal, 5, 123–130.

Darlington, G.J. et al. (1980) Expression of liver phenotypes in cultured mouse hepatoma cells. J Natl Cancer Inst, 64, 809–819.

Denison, M.S. et al. (2011) Exactly the same but different: promiscuity and diversity in the molecular mechanisms of action of the aryl hydrocarbon (dioxin) receptor. Toxicol. Sci., 124, 1–22.

Denison, M.S. et al. (1988) Inducible, receptor-dependent protein-DNA interactions at a dioxin-responsive transcriptional enhancer. Proc Natl Acad Sci USA, 85, 2528–2532.

Dere, E. et al. (2011) Integration of genome-wide computation DRE search, AhR ChIP-chip and gene expression analyses of TCDD-elicited responses in the mouse liver. BMC Genomics, 12, 365.

Diani-Moore, S. et al. (2010) Identification of the aryl hydrocarbon receptor target gene TiPARP as a mediator of suppression of hepatic gluconeogenesis by 2,3,7,8-tetrachlorodibenzo-p-dioxin and of nicotinamide as a corrective agent for this effect. J. Biol. Chem., 285, 38801–38810.

Dobin, A. et al. (2013) STAR: ultrafast universal RNA-seq aligner. Bioinformatics, 29, 15–21.

Dougherty, E.J. and Pollenz, R.S. (2008) Analysis of Ah receptor-ARNT and Ah receptor-ARNT2 complexes in vitro and in cell culture. Toxicol. Sci., 103, 191–206.

Ernst, J. and Kellis, M. (2017) Chromatin-state discovery and genome annotation with ChromHMM. Nat. Protoc., 12, 2478–2492.

Ernst, J. and Kellis, M. (2012) ChromHMM: automating chromatin-state discovery and characterization. Nat. Methods, 9, 215–216.

Fader, K.A. et al. (2019) 2,3,7,8-Tetrachlorodibenzo-p-dioxin abolishes circadian regulation of hepatic metabolic activity in mice. Sci. Rep., 9, 6514.

Fader, K.A. et al. (2017) Convergence of hepcidin deficiency, systemic iron overloading, heme accumulation, and REV-ERBα/β activation in aryl hydrocarbon receptor-elicited hepatotoxicity. Toxicol. Appl. Pharmacol., 321, 1–17.

Giuliano, C.J. et al. (2019) Generating Single Cell-Derived Knockout Clones in Mammalian Cells with CRISPR/Cas9. Curr. Protoc. Mol. Biol., 128, e100.

Gorelik, A. et al. (2025) CRISPR screens and quantitative proteomics reveal remodeling of the aryl hydrocarbon receptor-driven proteome through PARP7 activity. Proc Natl Acad Sci USA, 122, e2424985122.

Hankinson, O. (2005) Role of coactivators in transcriptional activation by the aryl hydrocarbon receptor. Arch. Biochem. Biophys., 433, 379–386.

Heinz, S. et al. (2010) Simple combinations of lineage-determining transcription factors prime cis-regulatory elements required for macrophage and B cell identities. Mol. Cell, 38, 576–589.

Hirose, K. et al. (1996) cDNA cloning and tissue-specific expression of a novel basic helix-loop-helix/PAS factor (Arnt2) with close sequence similarity to the aryl hydrocarbon receptor nuclear translocator (Arnt). Mol. Cell. Biol., 16, 1706–1713.

Hogenesch, J.B. et al. (1998) The basic-helix-loop-helix-PAS orphan MOP3 forms transcriptionally active complexes with circadian and hypoxia factors. Proc Natl Acad Sci USA, 95, 5474–5479.

Huang, G. and Elferink, C.J. (2012) A novel nonconsensus xenobiotic response element capable of mediating aryl hydrocarbon receptor-dependent gene expression. Mol. Pharmacol., 81, 338–347.

Humphrey-Johnson, A. et al. (2015) Stability of the aryl hydrocarbon receptor and its regulated genes in the low activity variant of Hepa-1 cell line. Toxicol. Lett., 233, 59–67.

Jackson, D.P. et al. (2014) Ah receptor-mediated suppression of liver regeneration through NC-XRE-driven p21Cip1 expression. Mol. Pharmacol., 85, 533–541.

Joshi, A.D. et al. (2015) Homocitrullination is a novel histone H1 epigenetic mark dependent on aryl hydrocarbon receptor recruitment of carbamoyl phosphate synthase 1. J. Biol. Chem., 290, 27767–27778.

J, L. (2024) eulerr: Area-Proportional Euler and Venn Diagrams with Ellipses R Package.

Keith, B. et al. (2001) Targeted mutation of the murine arylhydrocarbon receptor nuclear translocator 2 (Arnt2) gene reveals partial redundancy with Arnt. Proc Natl Acad Sci USA, 98, 6692–6697.

Klein-Hitpass, L. et al. (1988) Synergism of closely adjacent estrogen-responsive elements increases their regulatory potential. J. Mol. Biol., 201, 537–544.

Klinge, C.M. et al. (2000) The aryl hydrocarbon receptor interacts with estrogen receptor alpha and orphan receptors COUP-TFI and ERRalpha1. Arch. Biochem. Biophys., 373, 163–174.

Kobayashi, A. et al. (1997) CBP/p300 functions as a possible transcriptional coactivator of Ah receptor nuclear translocator (Arnt). J. Biochem., 122, 703–710.

Krueger, F. (2015) Trim Galore!: A wrapper around Cutadapt and FastQC to consistently apply adapter and quality trimming to FastQ files, with extra functionality for RRBS data. Babraham Institute.

Kume, K. et al. (1999) mCRY1 and mCRY2 are essential components of the negative limb of the circadian clock feedback loop. Cell, 98, 193–205.

Labun, K. et al. (2019) CHOPCHOP v3: expanding the CRISPR web toolbox beyond genome editing. Nucleic Acids Res., 47, W171–W174.

Langmead, B. and Salzberg, S.L. (2012) Fast gapped-read alignment with Bowtie 2. Nat. Methods, 9, 357–359.

Layer, R.M. et al. (2018) GIGGLE: a search engine for large-scale integrated genome analysis. Nat. Methods, 15, 123–126.

Liao, Y. et al. (2014) featureCounts: an efficient general purpose program for assigning sequence reads to genomic features. Bioinformatics, 30, 923–930.

Lo, R. and Matthews, J. (2012) High-resolution genome-wide mapping of AHR and ARNT binding sites by ChIP-Seq. Toxicol. Sci., 130, 349–361.

Madak-Erdogan, Z. and Katzenellenbogen, B.S. (2012) Aryl hydrocarbon receptor modulation of estrogen receptor α-mediated gene regulation by a multimeric chromatin complex involving the two receptors and the coregulator RIP140. Toxicol. Sci., 125, 401–411.

Marlowe, J.L. et al. (2004) The aryl hydrocarbon receptor displaces p300 from E2F-dependent promoters and represses S phase-specific gene expression. J. Biol. Chem., 279, 29013–29022.

Ma, Q. et al. (2001) TCDD-inducible poly(ADP-ribose) polymerase: a novel response to 2,3,7,8-tetrachlorodibenzo-p-dioxin. Biochem. Biophys. Res. Commun., 289, 499–506.

Mei, S. et al. (2017) Cistrome Data Browser: a data portal for ChIP-Seq and chromatin accessibility data in human and mouse. Nucleic Acids Res., 45, D658–D662.

Mouse ENCODE Consortium et al. (2012) An encyclopedia of mouse DNA elements (Mouse ENCODE). Genome Biol., 13, 418.

Müller-Dott, S. et al. (2023) Expanding the coverage of regulons from high-confidence prior knowledge for accurate estimation of transcription factor activities. Nucleic Acids Res., 51, 10934–10949.

Nault, R. et al. (2018) Comparison of Hepatic NRF2 and Aryl Hydrocarbon Receptor Binding in 2,3,7,8-Tetrachlorodibenzo-p-dioxin-Treated Mice Demonstrates NRF2-Independent PKM2 Induction. Mol. Pharmacol., 94, 876–884.

Nault, R. et al. (2016) Pyruvate Kinase Isoform Switching and Hepatic Metabolic Reprogramming by the Environmental Contaminant 2,3,7,8-Tetrachlorodibenzo-p-Dioxin. Toxicol. Sci., 149, 358–371.

Nault, R. et al. (2015) RNA-Seq versus oligonucleotide array assessment of dose-dependent TCDD-elicited hepatic gene expression in mice. BMC Genomics, 16, 373.

Quinlan, A.R. and Hall, I.M. (2010) BEDTools: a flexible suite of utilities for comparing genomic features. Bioinformatics, 26, 841–842.

Reyes, H. et al. (1992) Identification of the Ah receptor nuclear translocator protein (Arnt) as a component of the DNA binding form of the Ah receptor. Science, 256, 1193–1195.

Risso, D. et al. (2014) Normalization of RNA-seq data using factor analysis of control genes or samples. Nat. Biotechnol., 32, 896–902.

Robinson, M.D. et al. (2010) edgeR: a Bioconductor package for differential expression analysis of digital gene expression data. Bioinformatics, 26, 139–140.

Rowatt, A.J. et al. (2003) ARNT gene multiplicity in amphibians: characterization of ARNT2 from the frog Xenopus laevis. J. Exp. Zool. B Mol. Dev. Evol., 300, 48–57.

RStudio Team (2025) RStudio: Integrated Development for R RStudio, Inc., Boston, MA.

Subramanian, A. et al. (2005) Gene set enrichment analysis: a knowledge-based approach for interpreting genome-wide expression profiles. Proc Natl Acad Sci USA, 102, 15545–15550.

Takumi, T., Taguchi, K., et al. (1998) A light-independent oscillatory gene mPer3 in mouse SCN and OVLT. EMBO J., 17, 4753–4759.

Takumi, T., Matsubara, C., et al. (1998) A new mammalian period gene predominantly expressed in the suprachiasmatic nucleus. Genes Cells, 3, 167–176.

Tanguay, R.L. et al. (1999) Cloning and characterization of the zebrafish (Danio rerio) aryl hydrocarbon receptor. Biochim. Biophys. Acta, 1444, 35–48.

Tanguay, R.L. et al. (2000) Identification and expression of alternatively spliced aryl hydrocarbon nuclear translocator 2 (ARNT2) cDNAs from zebrafish with distinct functions. Biochim. Biophys. Acta, 1494, 117–128.

Tennant, B.R. et al. (2013) Identification and analysis of murine pancreatic islet enhancers. Diabetologia, 56, 542–552.

van der Velde, A. et al. (2021) Annotation of chromatin states in 66 complete mouse epigenomes during development. Commun. Biol., 4, 239.

Vogel, C.F.A. et al. (2007) RelB, a new partner of aryl hydrocarbon receptor-mediated transcription. Mol. Endocrinol., 21, 2941–2955.

Westermann, L. et al. (2022) Wildtype heterogeneity contributes to clonal variability in genome edited cells. Sci. Rep., 12, 18211.

Wilson, S.R. et al. (2013) The tumor suppressor Kruppel-like factor 6 is a novel aryl hydrocarbon receptor DNA binding partner. J. Pharmacol. Exp. Ther., 345, 419–429.

Xu, C.-X. et al. (2010) Disruption of CLOCK-BMAL1 transcriptional activity is responsible for aryl hydrocarbon receptor-mediated regulation of Period1 gene. Toxicol. Sci., 115, 98–108.

Yao, E.F. and Denison, M.S. (1992) DNA sequence determinants for binding of transformed Ah receptor to a dioxin-responsive enhancer. Biochemistry, 31, 5060–5067.

Zhang, Y. et al. (2008) Model-based analysis of ChIP-Seq (MACS). Genome Biol., 9, R137.

Zheng, R. et al. (2019) Cistrome Data Browser: expanded datasets and new tools for gene regulatory analysis. Nucleic Acids Res., 47, D729–D735.

